# A transcriptomic atlas of *Aedes aegypti* reveals detailed functional organization of major body parts and gut regional specializations in sugar-fed and blood-fed adult females

**DOI:** 10.1101/2021.12.19.473372

**Authors:** Bretta Hixson, Xiao-Li Bing, Xiaowei Yang, Alessandro Bonfini, Peter Nagy, Nicolas Buchon

## Abstract

Mosquito vectors transmit numerous pathogens, but large gaps remain in our understanding of their physiology. To facilitate future explorations of mosquito biology, with specific attention to the major vector *Aedes aegypti*, we have created Aegypti-Atlas (http://aegyptiatlas.buchonlab.com/), an online resource hosting RNAseq profiles of *Ae. Aegypti* body parts (head, thorax, abdomen, gut, Malpighian tubules, and ovaries), gut regions (crop, proventriculus, anterior and posterior midgut, and hindgut), and a time course of blood meal digestion in the gut. Using Aegypti-Atlas, we provide new insights into the regionalization of gut function, blood feeding response, and immune defenses. We find that the anterior and posterior regions of the mosquito midgut possess clearly delineated digestive specializations which are preserved in the blood-fed state. Blood feeding initiates the sequential transcriptional induction and repression/depletion of multiple cohorts of peptidases throughout blood meal digestion. With respect to defense, immune signaling components, but not recognition or effector molecules, show enrichment in ovaries. Basal expression of antimicrobial peptides is dominated by two genes, holotricin and gambicin, that are expressed in the carcass and the digestive tissues, respectively, in a near mutually exclusive manner. In the midgut, gambicin and other immune effector genes are almost exclusively expressed in the anterior regions, while the posterior midgut exhibits the hallmarks of immune tolerance. Finally, in a cross-species comparison between the midguts of *Ae. aegypti* and *Anopheles gambiae*, we observe that regional digestive and immune specializations are closely conserved, indicating that our data may yield inferences that are broadly relevant to multiple mosquito vector species. We further demonstrate that the expression of orthologous genes is highly correlated, with the exception of a ‘species signature’ comprising a small number of highly/disparately expressed genes. With this work, we show the potential of Aegypti-Atlas to unlock a more complete understanding of mosquito biology.

## Introduction

Hematophagous mosquito vectors contribute substantially to the world’s disease burden through the transmission of *Plasmodium* parasites and arboviruses [1]. With the rise of insecticide resistance [2–4], and the expansion of mosquito species’ ranges potentiated by climate change [5], new and more efficient means of vector control are needed, grounded in a solid understanding of mosquito physiology.

The mosquito gut is of special interest to vector control, as it is central to the hematophagous lifestyle and serves as the first interface between a mosquito and the pathogens it transmits. In recent years, many groups have used microarrays and RNAseq to create transcriptomic profiles of different mosquito tissues, especially midguts, in the context of infection [6–13]. Here, we aimed to complement these efforts by (a) contextualizing the adult female gut’s transcriptome with additional profiles of the whole body and other major body parts and organs (head, thorax, abdomen, Malpighian tubules, ovaries) (b) exploring regionalization of gut function by profiling the anatomically distinct regions of the adult female gut (crop, proventriculus, anterior midgut, posterior midgut, hindgut) and (c) examining transcriptional responses to blood feeding in the anterior and posterior midgut and at multiple timepoints (6, 24, and 48 hrs) after blood meal ingestion. For this project, we opted to profile *Aedes aegypti*, as it is among the most important insect vectors [14], with an increasing worldwide range [15], and a recently updated genome assembly [16]. All the resulting transcriptional data can be accessed at the Aegypti-Atlas online database we constructed (http://aegyptiatlas.buchonlab.com/).

The FlyAtlas project [17, 18] contributed to the field of insect physiology, not only by characterizing the functions associated with the major body parts and organs of Drosophila, but by making it possible to quickly ascertain where in the body, and in what quantity, any gene of interest is transcribed. FlyAtlas also showed how the various contributions of different organs added up to constitute the whole fly transcriptome, providing an estimate of the relative transcriptional yield of the parts of the Drosophila body. Similar anatomical datasets have been created for several species of *Anopheles* [11,19,20], but not for *Aedes* mosquitoes. While transcriptional profiles have been created for various *Ae. aegypti* body parts in diverse studies (*e.g*. head [21], Malpighain tubules [22], midgut [7,9,23–27], ovaries [28, 29], carcass [28, 30]), the lack of uniformity of strain and methodology between these studies makes direct comparison difficult. We elected to profile the major body parts and organs of female *Ae. aegypti* in parallel with the gut with the dual goals of (a) creating an at-a-glance reference for the anatomical distribution of transcripts, genome-wide and (b) facilitating the identification of tissue-specific marker genes which may, in the future, be useful in the construction of tissue-specific expression systems.

Insect guts are highly regionalized organs, with specialized functions differentially distributed along their length [31–34]. The mosquito gut is divided into five anatomically distinct regions: foregut (comprising pharynx, dorsal diverticula, and crop), proventriculus, anterior midgut, posterior midgut, and hindgut. The crop and dorsal diverticula are muscular cuticle-lined sacs which store imbibed sugar for gradual release into the midgut [35]. In *Anopheles gambiae*, the proventriculus and anterior midgut were found to exhibit an enrichment of transcripts encoding antimicrobial peptides (AMPs) and anti-*Plasmodium* factors [13] suggesting that these regions serve a special defensive function against orally acquired pathogens. Within the midgut, digestive functions are believed to be divided between the anterior and posterior regions, with the former specializing in the digestion of nectar but playing no direct role in the digestion of the blood meal, which is processed in the posterior midgut [36]. The hindgut receives and eliminates excreta from the midgut and urine from the Malpighian tubules [35]. Genome-wide transcriptomic profiles have the potential to reveal much more detailed information about regionalized function in the mosquito gut. However, to our knowledge, only a single microarray-based study has ever characterized the transcriptomes of individual mosquito midgut regions [13] and no transcriptomic profiles have been generated for either foregut or hindgut tissues in any mosquito species. To address this lack, we created RNAseq profiles for all five regions of the *Ae. aegypti* gut. Additionally, to more thoroughly explore the roles of the anterior and posterior midgut in blood meal digestion, we created profiles for the two regions 24 hrs after ingestion of a blood meal.

The *Ae. aegypti* gut’s response to the ingestion of a blood meal has been characterized as a biphasic process. In the first phase mRNAs encoding a cohort of “early” serine endopeptidases, transcribed in response to juvenile hormone (JH) secretion during previtellogenic maturation, are translated [37–42]. In the second, ecdysone signaling mediates the transcription of a “late” cohort of endo and exopeptidases, with the transcripts of key peptidases and overall proteolytic activity typically peaking around 24 hrs post blood meal (pbm) [43–50]. The transcriptional timeline is, however, more complex than a simple early translation/late transcription paradigm might suggest. Several studies have, by RTqPCR and microarray, documented *de novo* transcription of peptidases and other genes peaking much earlier than 24 hrs pbm [51–53]. To better characterize the transcriptome of the gut, genome-wide, over the course of the digestion of a blood meal, we created profiles for guts at 6, 24, and 48 hrs pbm to compare with a sugar-fed baseline.

With RNAseq profiles for body parts, gut regions, and blood-fed guts in hand, we launched a broad investigation into multiple aspects of mosquito biology. Our inquiries were guided by the following questions: What functions are associated with each body part? What are the functional specializations of the gut regions? How does the gut transcriptome change in the context of blood feeding, and how are those changes distributed in the midgut? What can our atlas tell us about the organization of mosquitoes’ antimicrobial defenses? Finally, in a cross-species comparison between the midguts of *Ae. aegypti* and *Anopheles gambiae*, we asked whether the gut’s structure-function relationship is conserved between two hematophagous mosquito vector species.

## Results

### The transcriptional landscape of an adult mosquito reflects the embryonic origins and functional signatures of its major tissues

In order to develop a transcriptomic atlas of some of the main body parts of *Ae. aegypti*, we generated RNAseq profiles from the whole bodies and dissected body parts (head, thorax, abdomen, gut, Malpighian tubules, and ovaries) of sucrose-fed adult mated female mosquitoes of a field-derived ‘Thai’ strain from a region where arbovirus transmission is endemic [54]. For the dissected body parts (but not whole body) we omitted the anterior-most section of the thorax as well as the final segments of the abdomen to exclude transcripts from salivary glands and from sperm in the spermatheca (see S1.1A for diagram). We first evaluated the relative variance between body part and whole-body profiles by Principal Component Analysis (PCA) (Fig 1A). All transcriptomes from the same body part clustered together closely, providing evidence for the replicability of our data. On PC1, body parts were grouped in a manner that accorded with their divergent embryonic origin (primarily mesoderm/ectoderm for head, thorax, and abdomen, endoderm for gut and Malpighian tubules) and whole-body transcriptomes were clustered in between. Ovaries, which contain the germline, strongly separated from the other, exclusively somatic, body parts on PC2. Altogether, PCA indicated that the transcriptomes of body parts are largely defined by their embryonic origin, and that germline tissues possess a highly distinctive transcriptional signature.

**Fig 1.**
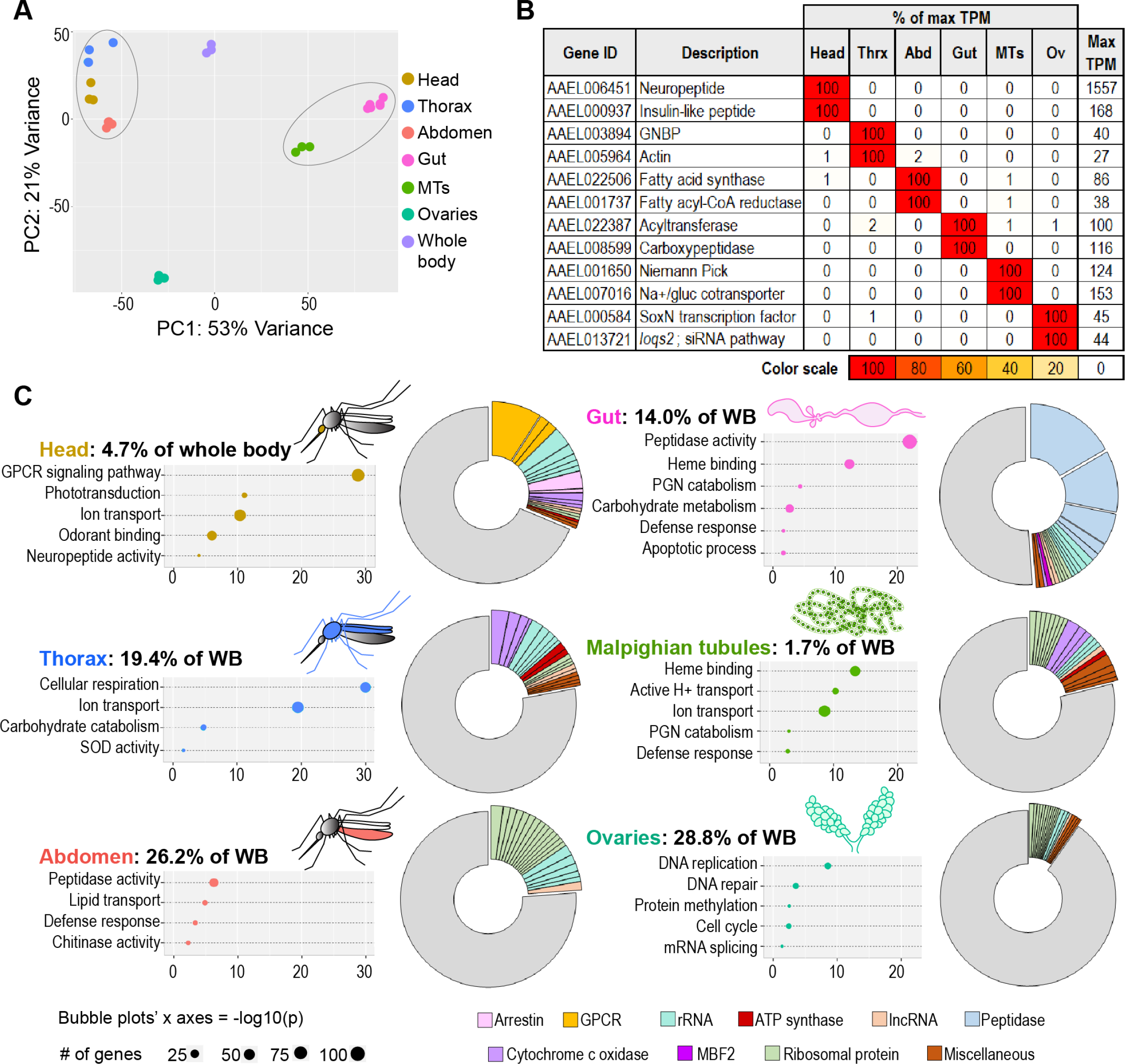
The transcriptomes of *Aedes aegypti* body parts reflect their embryonic origins and functional specializations. (A) PCA of the transcriptomes of whole body and body parts; 3 to 6 replicates per body part. Circled clusters contain body parts of dominantly endodermal (left) and mesodermal/ectodermal (right) derivation. (B) Body part-specific putative tissue markers; expression values are scaled as a percentage of the maximum expression observed for each gene. (C) Calculated estimates of body parts’ contributions to whole body transcriptome; Bubble plots: TopGO GOEA of genes enriched 5x (head, gut thorax, Malpighian tubules, abdomen) or 2x (ovaries) relative to whole body (DESeq2, padj <0.05); Pie charts: 20 highest expressed genes per body part, grouped by category/function.

Each body part we sequenced expressed a set of unique or nearly unique transcripts some of which, we hypothesize, may be exclusive tissue markers. We conducted a census of putative tissue markers (here defined as genes expressed at five or more transcripts per million (TPM) in one body part and enriched at least fifty-fold compared to all other body parts). Figure 1B presents a selection of markers from each body part (full census in S1 Table). The quantity of markers ranged from 8 in the abdomen to 391 in the ovaries, which expressed more markers than all other body parts combined; nearly one in fifty of all genes in the genome qualified as an ovary marker. We further noted that ovaries simultaneously expressed the greatest number of highly enriched transcripts *relative to other dissected body parts* (S1.1B Fig) and the smallest number of highly enriched transcripts *relative to whole body* (S1.1C Fig). This apparent paradox led us to hypothesize that the ovaries make a disproportionately large contribution to the transcriptome of the whole female mosquito. A gene that is uniquely expressed in the ovaries will not be highly enriched relative to the whole mosquito body if the ovarian transcriptome comprises a large proportion of the whole body’s transcriptome.

To validate our sequencing results, we performed RT-qPCR amplification of one putative marker gene from each body part. We selected a translation factor (AAEL013144) as a reference gene for our assay over more traditional candidates [55], on the basis of its robust expression and relatively small variation across samples (S1.2A Fig). The results of our RT-qPCR confirmed the anatomical specificity of each selected marker (S1.2B Fig).

The existence of transcripts specific to each body part allowed us to calculate their relative contribution to the whole body’s transcriptome (see methods). Expressed as percentages, the body part contributions we obtained were: head (4.7%), thorax (19.4%), abdomen (26.2%), gut (14.0%), Malpighian tubules (1.7%), and ovaries (28.8%). Collectively, these scaling factors sum to 94.9%. We validated our estimates by using them to predict the whole-body expression of every gene in the genome, then plotting those predictions against observed values from the whole-body transcriptome (S1.3A Fig). Calculated and observed values were highly correlated (slope of 1.00, R^2^=0.95) after removal of a single outlying gene (AAEL018689, a mitochondrial ribosomal RNA) which was underpredicted by our calculation. This analysis validates our hypothesis that the ovaries contribute a disproportionate number of transcripts to the whole-body transcriptome. Further, our data demonstrate that, due to the preponderance of ovarian contribution, multiple tissues have only a low representation in the body, severely limiting the ability to detect tissue-specific changes in whole-mosquito experiments.

To examine the biological processes and molecular functions enriched in each body part, we performed a Gene Ontology Enrichment Analysis (GOEA) of genes 5x enriched in each body part (in comparison to whole body, DESeq2 padj <0.05). An extended list of GO categories may be found in S2 Table. We complemented this approach by identifying the twenty highest expressed genes in each body part (pie charts in Fig 1C), reasoning that the highest expressed genes in a transcriptome may also give insight into function. In brief, the head showed enrichment for sensory machinery and neuronal signaling, the thoracic carcass (which houses flight muscle) showed signs of enhanced metabolic activity, and the abdominal carcass (which contains mostly fat body) displayed enrichment for regulatory proteolytic enzymes (CLIP-domain serine endopeptidases), defense, and lipid transport. The gut was enriched with defense-related genes including immune-activating peptidoglycan recognition proteins (PGRPs), AMPs, and immune-modulating amidase PGRPs. It also showed enrichment for digestive enzymes, especially peptidases. Notably, its three most prevalent transcripts were the non-CLIP serine endopeptidases early trypsin (*ET*), female-specific chymotrypsin (*CHYMO*), and juvenile hormone-regulated chymotrypsin (*JHA15*) which, together, accounted for more than one third of its transcriptome. (Note: Vectorbase gene IDs for all genes named in this text are listed in S3 Table). The Malpighian tubules, which serve a function analogous to mammalian kidneys, expressed large quantities of ion transporters, especially vacuolar ATPases, which may be required to create proton gradients for use in secondary active transport [56]. Because the ovaries did not express many highly enriched genes, we adjusted our analysis to examine genes expressed 2x higher than in the whole body and discovered categories related to cell division and germline maintenance. Overall, we concluded that the most prominent functions in each body part were consistent with expectations for the tissues they house. We take these results as a validation of our data set, and as confirmation that the segments of the mosquito carcass (head, thorax, abdomen) perform well as proxies for important tissues (brain/eyes, flight muscle, fat body).

### Gut region-specific transcriptomes reveal functional compartmentalization of the mosquito midgut

To evaluate the regionalization of gut function, we generated RNAseq profiles for the five main regions of the mosquito digestive tract: crop and dorsal diverticula (hereafter referred to as “crop” for brevity), proventriculus, anterior midgut, posterior midgut, and hindgut. A PCA of these profiles (Fig 2A) demonstrated clear segregation between regions, and close clustering of all replicates from the same region, indicating clearly distinct transcriptomes. The anterior and posterior midgut, which are predominantly derived from endoderm, clustered together opposite the crop and hindgut, which are of ectodermal origin. The proventriculus, which is partially derived from both germ layers, was intermediate between these two clusters on PC1. Whole-gut transcriptomes clustered closely with posterior midgut transcriptomes, suggesting that the posterior midgut contributes more to the transcriptome of the whole gut than do the other regions. Accordingly, very few genes in the posterior midgut showed enrichment compared to the whole gut (S2A Fig). GOEA on genes 5x enriched in each region compared to the whole gut (Fig 2B), complemented by an examination of the twenty highest expressed genes in each gut region (Fig 2C) revealed that the ectodermally-derived regions of the gut (crop and hindgut) expressed genes involved in chitin metabolism, lipid metabolism, ion transport and numerous CLIP-domain serine endopeptidases. The proventriculus was notably enriched for wingless signaling and defensive genes (lysozymes, and the highly expressed AMP gambicin, *GAM1*). In the anterior midgut, carbohydrate metabolism and heme-binding proteins (predominantly cytochrome P450s) were the most prominently enriched categories. The highest expressed gene in this region was a member of the MBF2 family of transcription factors of unknown function in mosquitoes. As the posterior midgut transcriptome closely resembled that of the whole gut, only a handful of genes were enriched 5x in this region relative to whole gut. We also noted that a large proportion of the genome was either unexpressed or little-expressed in the posterior midgut relative to the other gut regions (S2B Fig). To better identify genes enriched in the posterior midgut, we calculated enrichment against an aggregate of the other four gut regions. We found that the posterior midgut is highly enriched for peptidases (especially non-CLIP serine endopeptidases) and genes involved in translation. Other enriched categories include SRP-dependent targeting, amino acid transport and amino acid metabolism. Together these categories describe a region that is primed to translate and secrete large quantities of peptidases, and to process and absorb the resulting free amino acids. Overall, our data demonstrate that the five gut regions are highly specialized functional units.

**Fig 2:**
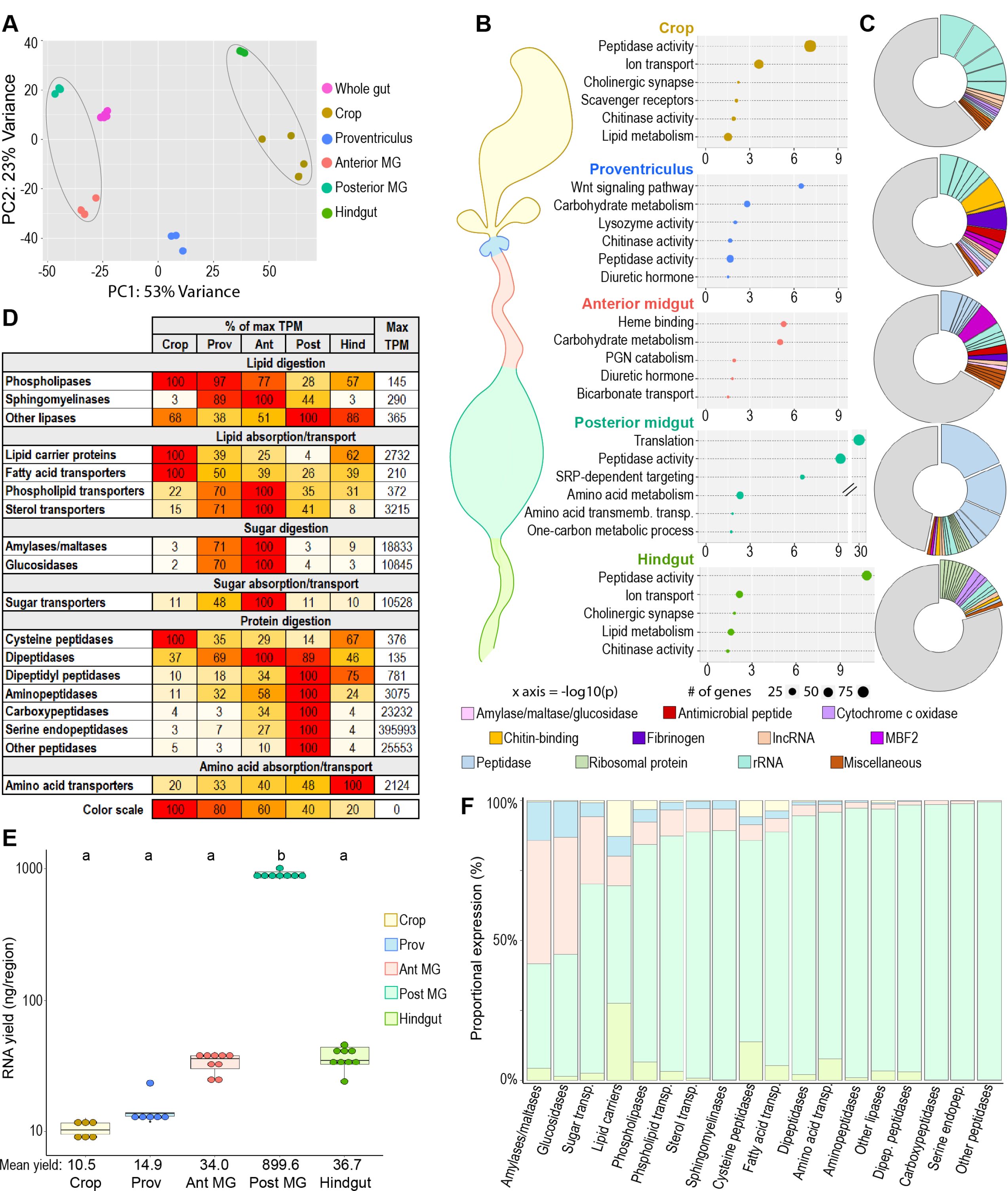
Mosquito gut regions’ transcriptomic investments and outputs reveal digestive specializations. (A) Principal component analysis of the transcriptomes of whole gut and gut regions; 3 to 6 replicates. Circled clusters contain body parts of partial endodermal (left) and ectodermal (right) derivation. (B) Bubble plots: TopGO GOEA of regionally enriched genes (5x over whole gut for crop, proventriculus, and anterior midgut, 5x over other regions for posterior midgut) in *Aedes aegypti* (DESeq2, padj <0.05) (C) Pie charts: 20 highest expressed genes per region, grouped by category/function. (D) Regional investments in categories of digestive enzymes and transporters, scaled as a percentage of the most invested region (as measured by cumulative TPM). (E) Estimates of regions’ transcriptional output by empirical quantification of RNA yield; statistics: One-way ANOVA, Tukey HSD. (F) Estimated output of digestive enzymes/transporters by region (categorical investments weighted by regional RNA yield).

The central function of the gut is digestion, a highly sequential process. To better understand how digestion and nutrient absorption are distributed along the gut, we examined the regional cumulative expression of enzymes and transporters putatively involved in the digestion and absorption of lipids, carbohydrates, and proteins/amino acids. We postulated that the cumulative expression of a given set of digestive enzymes/transporters of common function reflects the *investment* of each region in that function (what fraction of their transcriptome they dedicate to that process). We first assembled lists of lipases, lipid-transporting proteins, amylases/maltases, glucosidases, sugar transporters, peptidases, and amino acid transporters with a probable digestive function (excluding genes with anticipated non-digestive function, *e.g.*, peptidases annotated as, or orthologous with, CLIP-domain serine endopeptidases, caspases, proteasome components, *etc.*). For complete lists and inclusion criteria, see S4 Table. We summed the expression of the genes from each category (in TPM) and calculated their relative expression by region (Fig 2D). The highest investments in different categories of lipases were divided among several regions, with phospholipases receiving the greatest investment from crop and anterior midgut, sphingomyelinases from anterior midgut, and the remainder of lipases modestly more expressed in the posterior midgut and hindgut. The crop was the greatest investor in lipid carrier proteins and fatty acid transporters, while the anterior midgut showed the greatest investment in phospholipid and sterol transporters. The highest investments in sugar digestion/absorption and protein digestion were found in the anterior and posterior midgut, respectively. The hindgut was a disproportionate investor in amino acid transporters, suggesting that many of the products of protein digestion from the midgut are absorbed there. These data demonstrate that distinct gut regions invest different amounts of their respective transcriptomes in the digestion and absorption of different nutrients, hinting at a system of specialization/sequential processing of ingested materials.

Caution should be used in the interpretation of cumulative expression data, as *investment* (in TPM) is distinct from *output* (*i.e.*, the number of transcripts of a given gene or category produced in a region). The five regions do not produce equal numbers of transcripts overall, so their transcriptional investments must be scaled by their total transcriptional *yield* to achieve an estimate of their relative contributions to specific functions. We measured the RNA content from each region (Fig 2E) and scaled the cumulative expression values from Fig 2D by yield to gain an estimate of the transcriptional output of each region with respect to each digestive category (Fig 2F). We found that the posterior midgut, by virtue of its outsized transcriptional yield, is the dominant contributor not only of peptidases, but of most categories of transcripts with digestive and absorptive functions. However, the proventriculus and anterior midgut together are the source of more than half of all amylases/maltases and glucosidases and contribute a sizeable minority of all sugar transporters. These observations cement our conclusion that the anterior midgut specializes in the digestion and absorption of carbohydrates, while the posterior midgut is responsible for most protein and lipid digestion.

### Blood feeding initiates a series of transcriptomic shifts over the course of digestion

The gut of the hematophagous mosquito undergoes dramatic changes upon blood feeding. To better understand these changes at a transcriptome-wide level, we generated RNAseq profiles for sugar-fed guts as well as guts at 4-6 hrs (hereafter referred to as 6 hrs for brevity), 24 hrs, and 48 hrs after feeding on a live chicken. PCA (Fig 3A) demonstrates that the transcriptome changes throughout at least the first 24 hrs pbm, before reverting to a near-basal condition by 48 hrs pbm, suggesting that the transcriptional regulation of digestion is a very dynamic process.

**Fig 3:**
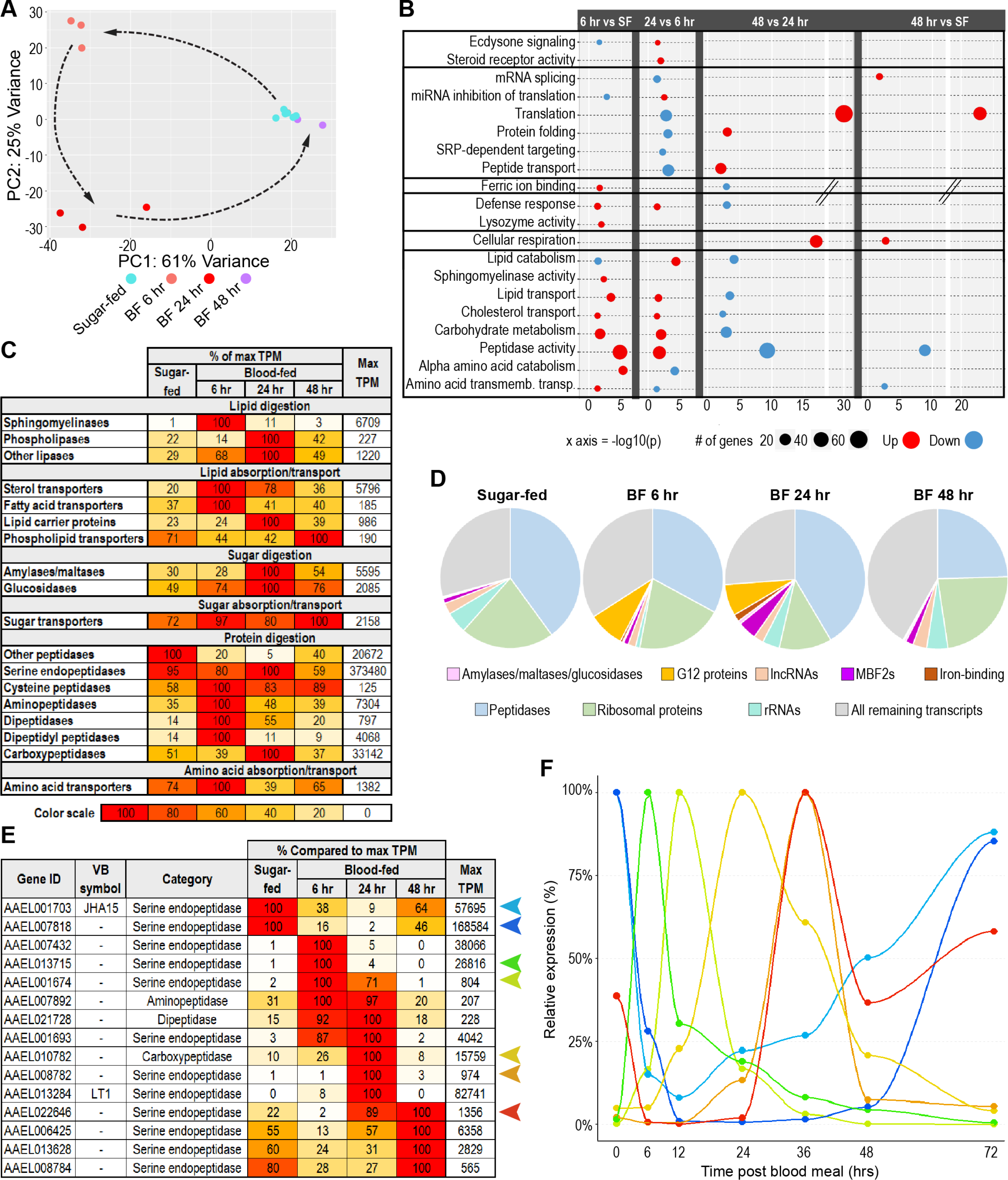
The gut’s transcriptome shifts continuously throughout blood meal digestion. (A) PCA of the transcriptomes of sugar-fed and blood-fed guts; 3 to 6 replicates per condition. (B) TopGO GOEA of up to 400 most upregulated and 400 most downregulated genes in each comparison (DESeq2, padj <0.05, minimum of 2x fold-change). (C) Investments in categories of digestive enzymes and transporters by timepoint/condition, scaled as a percentage of highest investment (as measured by cumulative TPM). (D) Proportional investment (out of whole transcriptome) in selected gene categories/families by timepoint/condition. (E) Expression of selected peptidases, scaled as a percentage of maximum expression. (F) Relative expression of selected peptidases at 0, 6, 12, 24, 36, 48, and 72 hrs post-blood meal measured by RT-qPCR; reference gene: AAEL009653; 3 replicates.

To evaluate the functional changes to the blood-fed gut’s transcriptome, we performed a GOEA (Fig 3B) comparing the gut at each time-point to its state at the time-point prior. Guts at the final timepoint (48 hrs) were also compared to sugar-fed guts. The analysis was limited to the 400 most responsive genes (DESeq2 padj < 0.05, minimum fold change of 2). We found that transcripts of *EcR*, the sole gene in the ecdysone signaling GO category, were significantly reduced in the gut at 6 hrs pbm, while at 24 hrs pbm, after the release of ecdysone peaks [57], not only did the level of *EcR* transcripts rebound but two other steroid receptors, *USP* (encoding the co-receptor to EcR) and *HR3*, were enriched. At 24 hrs pbm, multiple categories pertaining to mRNA processing, translation, protein folding, and protein secretion were significantly downregulated. Several of these categories had rebounded by 48 hrs pbm, and it was notable that translation was one of very few categories that was significantly enriched in 48-hr blood-fed guts compared to exclusively sugar-fed guts.

Several of the categories regulated by blood feeding appear to reflect necessary adaptations to the changing luminal environment. Six hours following blood meal ingestion, ferric iron-binding proteins (specifically two ferritin subunit precursors, AAEL004335 and AAEL007385) were upregulated and the lysozyme activity and defense response categories (comprising two lysozymes *LYSC11*, and *LYSC4* and the AMPs *DEFC*, *CECD*, and *CECN*) were also induced, in agreement with the previously described increase of ROS and bacterial density pbm [58]. At 24 hrs pbm, two additional defensins (*DEFA* and *DEFD*) were enriched, as well as two peptidoglycan-degrading amidases (*PGRP-SC1*, and *PGRPS4*), orthologs of which, in Drosophila, modulate the activation of the immune deficiency (IMD) pathway [59]. At 48 hrs pbm, both the ferric iron-binding and defense response categories were downregulated relative to 24 hrs, which is unsurprising as most of the luminal content had been evacuated by that time (S3.1A Fig). We also noted a significant induction of genes involved in cellular respiration which could reflect the metabolic expenditure required for the peristaltic evacuation of the blood bolus. GO categories related to the digestion and absorption of macronutrients were nearly all upregulated at 6 hrs pbm and upregulated again at 24 hrs but downregulated to return to baseline by 48 hrs. In summary, GOEA indicates that blood feeding regulates the transcription of genes involved in diverse functions, including digestion, absorption, translation, and defense.

To characterize the changes in digestion/absorption in more detail, we calculated the cumulative investment of the gut in each digestive category at each timepoint (Fig 3C). Overall, cumulative transcript analysis echoed the results of our GOEA. Both showed that the gut’s transcriptional investment in sphingomyelinases and amino acid transporters peaked at 6 hrs pbm, and that carbohydrate digestive enzymes generally gained ground until 24 hrs pbm but declined again at 48 hrs. Our granular cumulative expression analysis revealed that aminopeptidases, dipeptidases, and dipeptidyl peptidases reached peak investment at 6 hrs, and carboxypeptidases at 24 hrs. Remarkably, the gut’s investment in serine endopeptidases (non-CLIP), which constitute the majority of peptidase transcripts in the gut by nearly an order of magnitude, remained steady across sugar-fed, 6-hr, and 24-hr pbm guts, before plunging by approximately one third at 48 hrs pbm.

We extended our cumulative investment approach to other functions in the gut by aggregating transcripts of other transcriptionally prominent families and comparing their relative proportions across all timepoints (Fig 3D). This revealed the dramatic temporary increase in the expression of a family of twelve genes containing the insect allergen domain (IPR010629) which are orthologous to the *An. gambiae* G12 gene and have been found to possess hemolytic, cytolytic, and antiviral properties [60]. We also observed that the family of MBF2 transcription factors - one member of which was previously noted among the highest expressed genes of the anterior midgut (Fig 2C) - surged from just over 1% of the transcriptome to more than 4% at 24 hrs pbm, partially subsiding to approximately 1.7% by 48 hrs pbm. Our comparison of summed transcripts yielded an apparent paradox: while GOEA demonstrated the upregulation of multiple peptidases at 6 and 24 hrs pbm, and no significant downregulation of the category, the gut’s overall investment in peptidases (cumulative TPM) remained nearly constant in sugar-fed, 6-hr, and 24-hr pbm guts. This apparent stability is explained by the depletion of transcripts from the small “early” cohort of peptidases (*ET*, *CHYMO*, and *JHA15*) which compensated for the upregulation of other peptidases. It should be noted that, while peptidase *investment* remained flat, increased RNA yield (S3.1B Fig) caused peptidase transcript *output* to increase significantly at 24 hrs pbm (S3.1C Fig).

To more closely examine the dynamics of peptidase transcription in the blood-fed gut, we performed a clustering analysis (S3.2A Fig) which revealed that the transcript levels of many peptidases in the gut change dramatically over the course of blood meal digestion, that these changes come in the form of both induction and repression/depletion, frequently followed by a return to baseline at the subsequent timepoint, and that they manifest in multiple successive waves. Our clustering analysis divided the dynamically expressed gut peptidases into three waves of induced transcripts (rapid, intermediate, and delayed), and three corresponding waves of repressed or depleted transcripts. Among induced peptidases, the rapid and delayed waves comprise genes that peaked in our data set at 6 hrs and 24 hrs pbm, (23 and 40 genes respectively). Our clustering analysis implied that another, “intermediate” cohort of induced peptidases (16 genes) nests somewhere between these waves. These genes show sustained induction from 6 to 24 hrs, and likely peak at some time between these two timepoints. The waves of rapid, intermediate, and delayed repressed/depleted peptidases mirror the expression patterns of their induced counterparts in inverse. Figure 3E shows the expression of representative peptidases from each of the six cohorts across the blood feeding kinetic. We validated our findings by performing RT-qPCR on selected peptidases (two repressed/depleted and five induced genes) at 6, 12, 24, 36, 48, and 72 hrs pbm. We selected *RpS30* (AAEL009653), a housekeeping gene with robust and steady expression over the course of blood meal digestion, as a reference gene for our assay (S3.2B Fig). Figure 3F shows the relative expression of each peptidase (for unscaled data normalized only to the reference gene, see S3.2C Fig). At each timepoint we assayed, we captured one or more genes peaking sharply, further illustrating the dynamic nature of peptidase expression in the blood-fed gut. Most peptidases are organized in genomic clusters which, we hypothesized, could underlie co-regulation. We also examined the evolutionary relatedness of the peptidases from each temporal cluster by comparing their positions on a phylogenetic tree (composed using Geneious R11 software, Fig 3.3A). While sequence similarity corresponded closely with physical location in most cases, neither parameter was correlated with temporal clustering. Altogether, our data reveal that blood feeding initiates a well-orchestrated transcriptomic response in the gut, dominated by the dynamic up and down-regulation of peptidases throughout the entire process of digestion.

### Regional digestive specializations are preserved in the blood-fed midgut

Considering the extensive functional regionalization of the *Ae. aegypti* gut and the profound reshaping of the whole-gut transcriptome upon blood feeding, we asked whether regional differences are preserved in the blood-fed gut. To evaluate the regionalized effects of blood feeding, we generated transcriptomes for dissected anterior midguts (without proventriculus) and posterior midguts at 24 hrs pbm. PCA (Fig 4A) revealed that all profiles clustered both by region and by blood feeding status. As we previously noted (Fig 2A), there was little difference between sugar-fed whole gut and posterior midgut profiles. Upon blood feeding, both whole gut and posterior midgut shifted in tandem, maintaining proximity, indicating that blood feeding does not lessen the dominance of the posterior midgut’s transcriptional yield over that of the anterior. Both midgut regions shifted in response to blood feeding; moreover, they moved in the same directions with respect to PC1 and PC2, suggesting that the overall transcriptional responses of the two regions share some commonality. We noted that the relative investments of these two regions in most of the prominent functional categories did not change dramatically upon blood feeding (Fig 4B). However, two of the most remarkable changes in the transcriptome of blood-fed mosquitoes, the upregulation of MBF2 transcription factors and G12 genes (Fig 3D), were mainly localized to the posterior midgut.

**Fig 4:**
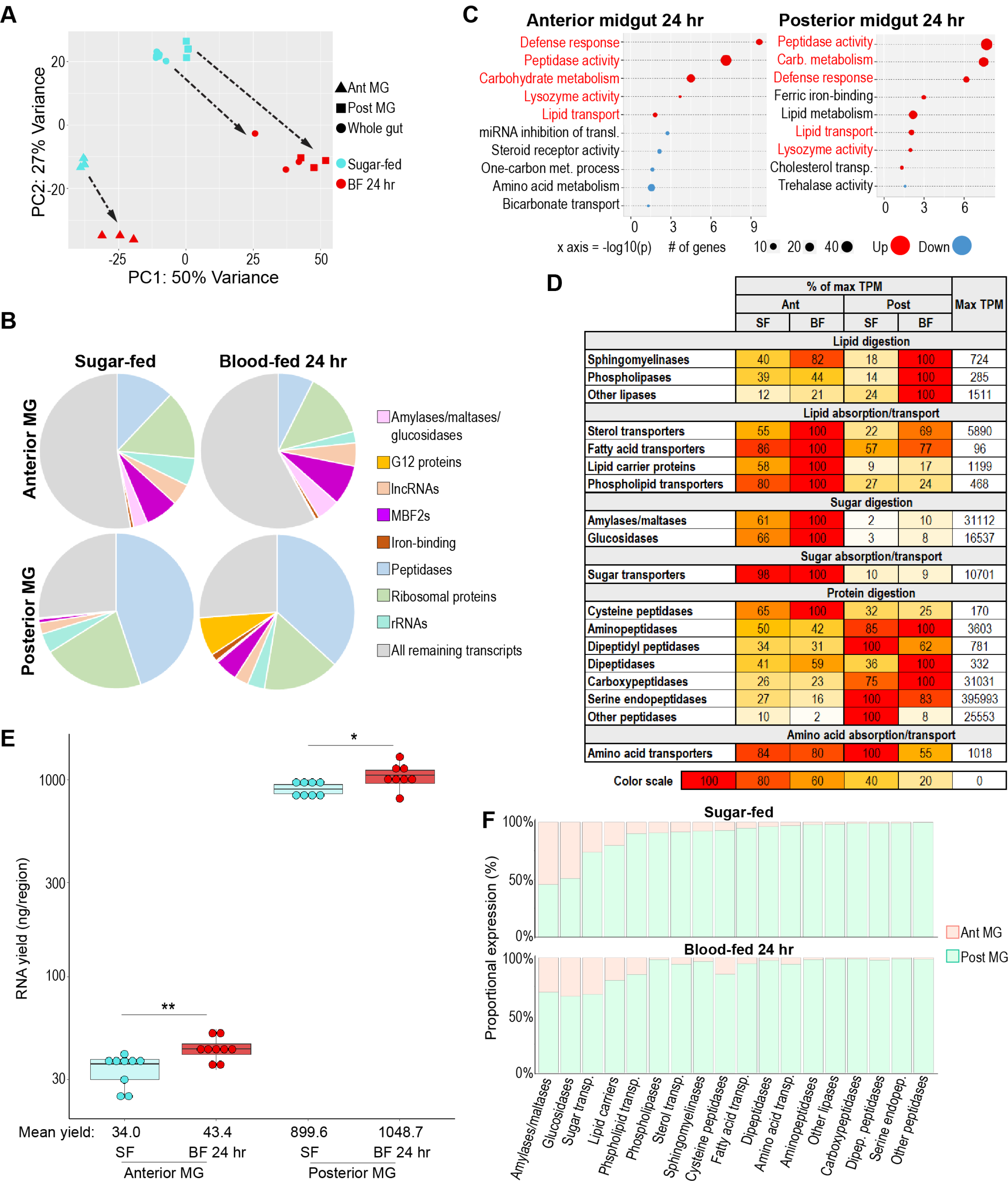
Regional specializations in sugar and protein digestion are preserved in the blood- fed midgut. (A) PCA of the transcriptomes of sugar-fed and 24-hr blood-fed whole guts and anterior and posterior midguts; 3 to 6 replicates per condition. (B) TopGO GOEA of up to 400 most upregulated and 400 most downregulated genes in blood-fed versus sugar-fed regions (DESeq2, padj <0.05, minimum of 2x fold-change inclusion criterion. (C) Proportional expression (out of whole transcriptome) of selected gene categories/families by region/condition. (D) Regional investments in categories of digestive enzymes and transporters, scaled as a percentage of the most invested region/condition (as measured by cumulative TPM). (E) Estimates of midgut regions’ transcriptional yield by empirical quantification of RNA in sugar-fed guts versus 24 hrs after feeding with an artificial blood replacement diet; statistics: unpaired t-test. (F) Estimated output of digestive enzymes/transporters by midgut region in sugar-fed and blood-fed mosquitoes (categorical investments weighted by regional RNA yield).

A GOEA of categories upregulated and downregulated by blood feeding in the anterior and posterior midgut (Fig 4C) mainly recapitulated changes already noted in blood-fed whole guts (Fig 3B). Most categories that were upregulated in one region were similarly upregulated in the other, albeit to differing extents. The fact that blood feeding promotes enrichment of many of the same digestive/absorptive categories in both anterior and posterior midgut could imply that the two regions are converging on a common role with respect to macronutrient exploitation. However, an analysis of cumulative expression (Fig 4D) demonstrated that the two regions’ respective investments in digestive enzymes and transporters remained highly divergent. Consistent with the results of the GOEA, the posterior midgut substantially increased its investment in lipase transcripts, with a smaller increase in the anterior midgut. By contrast, the anterior midgut kept a substantial edge in its investment in lipid transporters and lipid carrier proteins. The posterior midgut increased its investment in amylases/maltases and glucosidases by approximately five and two-fold, respectively, and the anterior midgut increased both by less than two-fold. However, the difference in baseline expression was such that the blood-fed anterior midgut’s investment was still ten times that of the blood-fed posterior midgut. Neither region substantially increased its investment in sugar transporters, peptidases, or amino acid transporters. When these regional investments were weighted by the increased RNA yield observed in both blood-fed midgut regions (Fig 4E), we found that the proportional output of the posterior midgut increased across most digestive categories, but that the anterior midgut still contributed a substantial minority of transcripts for sugar-digesting enzymes and sugar transporters (Fig 4F). Altogether, we concluded that both midgut regions participate in the transcriptional response to blood feeding, and that the blood-fed anterior and posterior midgut retain their regional specializations with respect to carbohydrate and protein digestion.

### Immune gene patterning reveals areas of immune activity and immune tolerance in the mosquito body and gut

As mosquitoes are prolific vectors of important human pathogens, their defensive functions are of paramount interest. We assembled a table recording the expression of immune genes in each category by body part, gut region, and blood-fed timepoint (S5 Table). Many genes associated with the siRNA and piRNA pathways display a strong tropism toward the ovaries (Fig 5A), possibly indicating that, in mosquitoes, siRNAs as well as piRNAs [61] are required to protect the germline from selfish elements. Accordingly, the gene *loqs2*, which has been described as an essential component of *Ae. aegypti’s* systemic siRNA response to Zika and dengue viruses [62] was restricted to the ovaries to such an extent that it qualified as an ovary-specific marker (Fig 1B). The ovaries were also the site of the highest expression of genes coding for components of Toll, IMD, and JAK-STAT signaling. Remarkably, genes coding for extracellular regulators (such as most CLIP-domain serine endopeptidases) and pattern recognition proteins (PGRPs and Gram-negative bacteria-binding proteins, GNBPs) did not show this pattern of expression. Instead, they were generally highly expressed in the head, thorax and abdomen, the three compartments that contain fat body. Even more remarkably, AMPs (Fig 5B) showed negligible expression in the ovaries. Our results suggest that, in the female germline, portions of the IMD, Toll and JAK-STAT pathways are uncoupled from upstream pattern recognition proteins and downstream AMP targets, possibly playing a regulatory role in some alternative process (*e.g.*, development).

**Fig 5.**
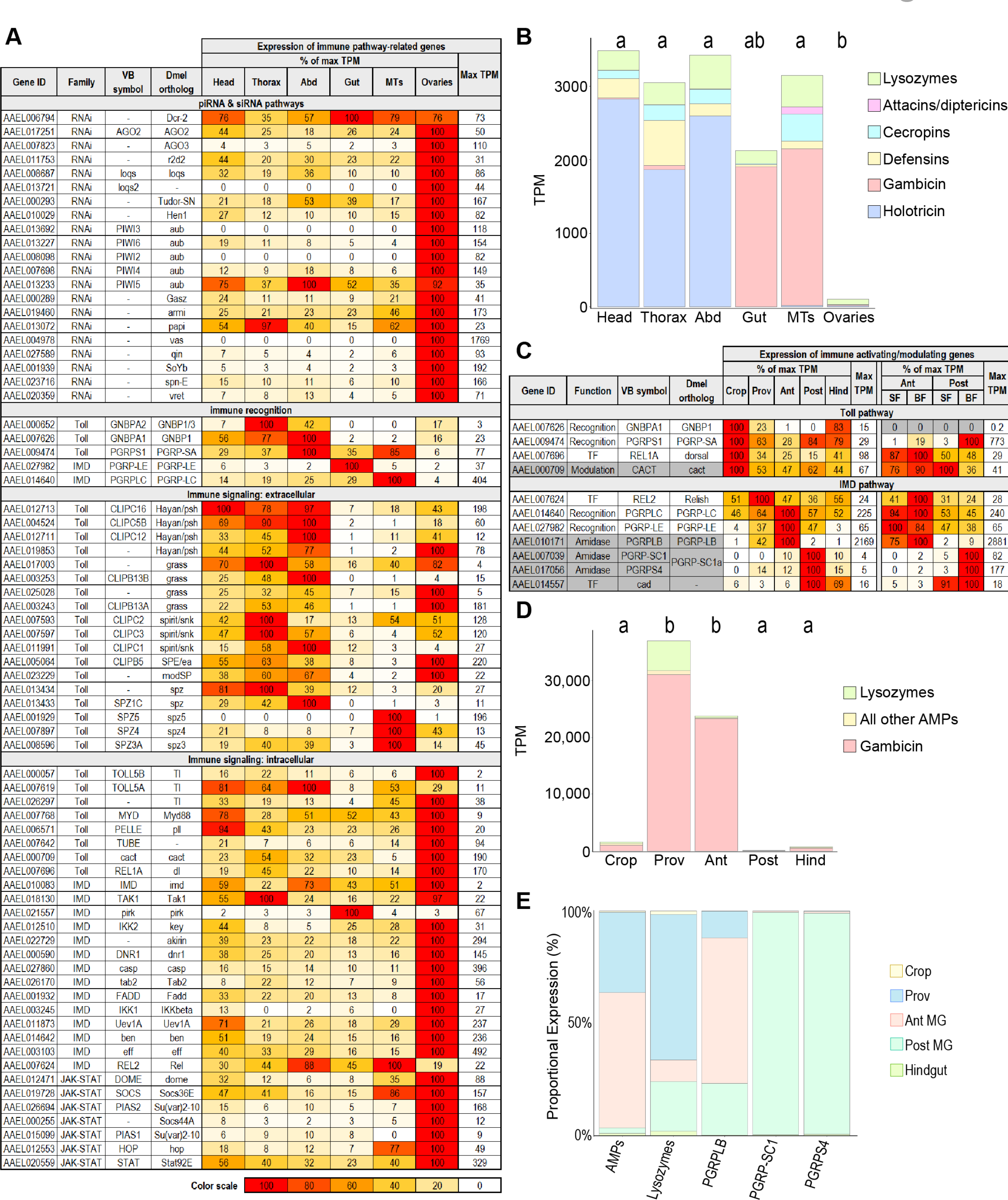
Immune gene patterning reveals zones of immune activity and tolerance in the mosquito body and gut. (A) Body parts’ investments in immune pathway-related genes, scaled as a percentage, scaled as a percentage of the most invested body part. (B) Cumulative antimicrobial peptide expression by body part (TPM); statistics: One-way ANOVA, Tukey HSD. (C) Expression of immune activating/modulating genes, scaled as a percentage of the highest expressing gut region. (D) Cumulative antimicrobial peptide expression by gut region (TPM); statistics: One-way ANOVA, Tukey HSD. (E) Output of immune-related genes and gene categories by gut region, weighted by regional RNA yield.

An examination of the distribution of AMPs and lysozymes in the body (Fig 5B) revealed that two AMPs, holotricin (*GRRP*) and gambicin, together accounted for most AMP transcripts in the mosquito body. These two AMPs were expressed in different body parts in a near mutually exclusive manner. The head, thorax, and abdomen of the mosquito predominantly expressed holotricin transcripts, with very little gambicin, while the gut and Malpighian tubules were dominated by gambicin expression and expressed very few holotricin transcripts.

Immune activity in the gut is tightly regulated as this organ interfaces with both commensal and pathogenic microbes. In Drosophila, the IMD pathway is the main pathway controlling AMPs in the midgut, and its activity is strongly constrained by expression of amidase PGRPs [59], while the Toll pathway is expressed mostly in the ectodermal portions of the gut. We found a similar pattern in the gut of *Ae. aegypti* (Fig 5C) with orthologs of Toll pathway recognition proteins most strongly expressed in the crop while IMD-activating PGRPs (*PGRPLC* and *PGRP-LE*) and IMD pathway components were enriched in the midgut. Investment in these IMD-activating PGRPs was highest in the anterior midgut, while immune-modulating PGRPs showed more divided expression. The amidase *PGRPLB* was most prominently expressed in the anterior midgut, but the short amidase PGRPs (*PGRP-SC1* and *PGRPS4*, orthologs of Drosophila *PGRP-SC1a* and *PGRP-SC1b*) were most strongly expressed in the posterior midgut. Likewise, an ortholog of caudal (*cad*), a transcription factor that limits AMP expression in the Drosophila midgut [63, 64], was profoundly enriched in the posterior midgut. Altogether, we found that the expression patterns of these key genes suggested enhanced immune vigilance in the anterior portion of the midgut, with hallmarks of immune tolerance prominent in the posterior midgut. We also observed that blood feeding enhanced the expression of the short amidase PGRPs in the posterior midgut, sharpening the regional divide.

A cumulative analysis of the expression of immune effectors (AMPs and lysozymes) in the gut showed that the proventriculus and anterior midgut, respectively, invest 3.6% and 2.3% of their transcriptomes in the expression of these effectors – especially gambicin (Fig 5D). The posterior midgut, by contrast, expressed just 110 TPM of AMPs and lysozymes combined, supporting our hypothesis that the posterior midgut is characterized by immune tolerance toward microbes. When AMP and lysozyme expression is weighted by gut regional RNA yield (Fig 5E) it is apparent that the anterior midgut and proventriculus contribute the majority of transcripts in these categories despite their relatively small transcriptional yields. Overall, the expression patterns of immune activators, modulators, and effectors suggests that the anterior portions of the midgut may exert strong selection of microbes upon entry, while the posterior midgut is a region of greater immune tolerance.

### Gut digestive and defensive regionalization is well conserved between the midguts of *Aedes aegypti* and *Anopheles gambiae*

*Ae. aegypti* and *An. gambiae* are both important hematophagous vectors, and their digestive tracts bear a clear anatomical similarity. To assess how similar these organs are at the transcriptomic level, we created RNAseq profiles for whole guts (comprising all regions from crop to hindgut) and midgut regions (proventriculus, anterior midgut and posterior midgut) from *An. gambiae* G3 mosquitoes and compared them to their *Ae. aegypti* counterparts. In a clustering analysis (Fig 6A) we found that the profiles of midgut regions and whole guts from each species were grouped similarly, with anterior midguts intermediate between proventriculi and posterior midguts. Also, in both species, whole guts clustered between anterior and posterior midgut regions but displayed more similarity to the latter, suggesting that for *An. gambiae*, as for *Ae. aegypti*, the transcriptional yield of the posterior midgut greatly exceeds that of the anterior.

**Fig 6.**
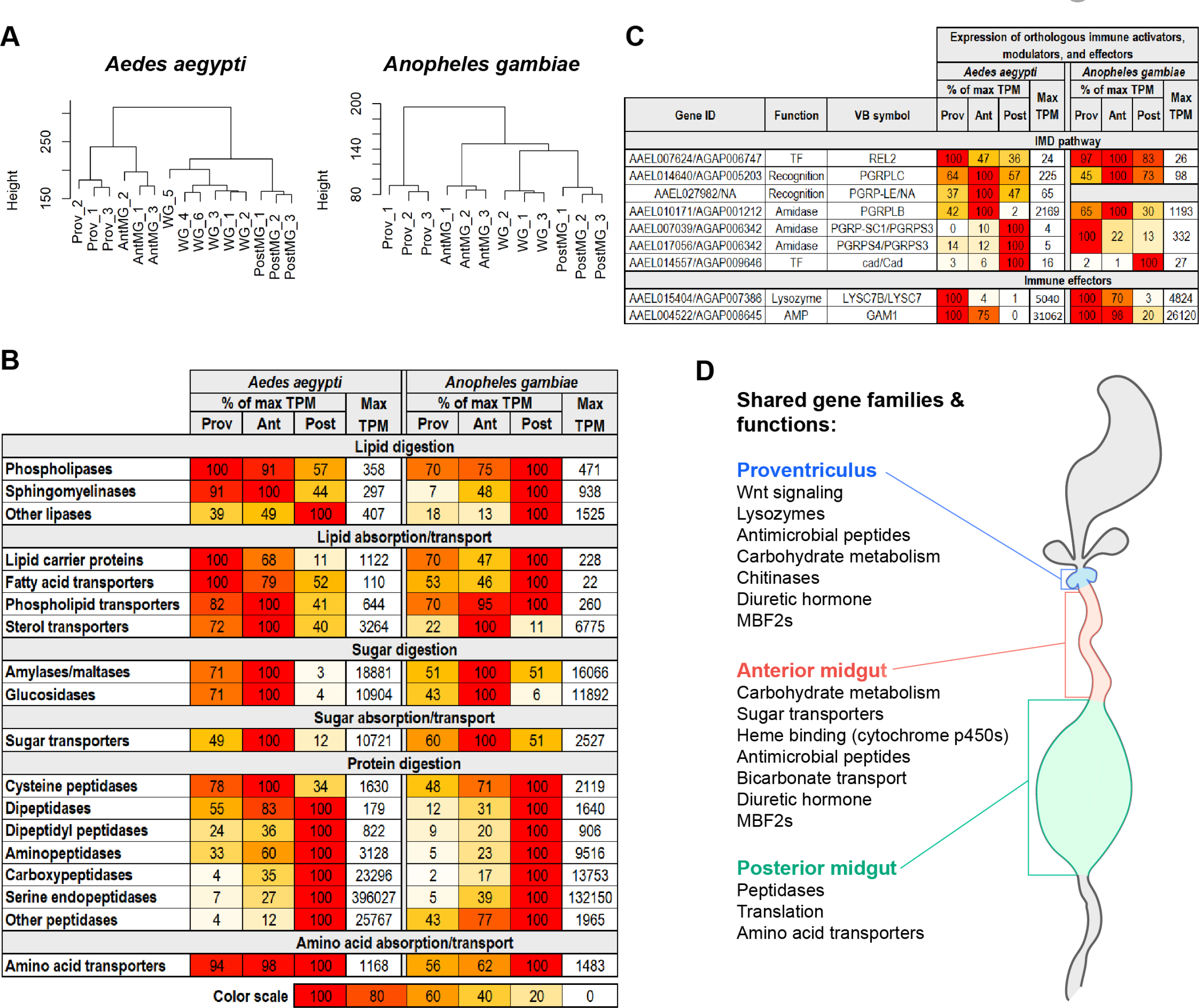
Digestive and defensive regional specializations are well conserved between the *Aedes aegypti* and *Anopheles gambiae* midguts. (A) Clustering analyses of the transcriptomes of *Aedes aegypti* and *Anopheles gambiae* whole gut and midgut regions; 3 to 6 replicates (B) Regional investments in categories of digestive enzymes and transporters, scaled as a percentage of expression in the most invested midgut region (as measured by cumulative TPM) (C) Expression of immune activating/modulating genes and effectors, scaled as a percentage of the expression in the most invested region. (D) Conserved regional enrichment of gene families and functions in *Ae. aegypti* and *An. gambiae* midgut regions, as evaluated by TopGO GOEA, investment analysis, and a comparison of the twenty highest expressed transcripts.

We repeated our analysis of regional transcriptomic investment in nutrient digestion and absorption by generating lists of genes encoding enzymes and transporters, in the same categories as in Figure 2C, summing their regional expression in TPM, and calculating their relative abundance. Since we lacked transcriptomes for whole body, crop, and hindgut for *An. gambiae*, it was necessary to adjust the inclusion criteria we employed (S4 Table) for our previous investment analysis. Complete lists of genes in each category with adjusted inclusion criteria can be found in S6 Table for *Ae. aegypti* and S7 Table for *An. gambiae.* Our comparison across the two species (Fig 6B) revealed overall conservation of gut structure-function. This was particularly evident for the anterior and posterior midguts which, respectively, remained the strongest investors in sugar and protein digestion/absorption. However, in *An. gambiae*, the posterior midgut appears to make a greater proportional investment in the digestion/absorption of both lipids and sugars. We also noted that, while the posterior midgut in both species was the dominant investor in peptidases, the amplitude of the *Ae. aegypti* investment far outstripped that of *An. gambiae* (S6A Fig), particularly with respect to serine endopeptidases.

To compare the distribution of defensive functions in the two midguts, we examined the expression patterns of important orthologous immune activators, modulators, and effectors between the two mosquitoes (Fig 6C) and found clear similarities. In both species, the IMD- activating *PGRPLC* receptor as well as the immune-modulating *PGRPLB* amidase were most strongly expressed in the anterior midguts, the transcription factor *cad* was most strongly expressed in the posterior midguts, and the orthologs of *GAM1* and *LYSC7B* (respectively the highest expressed AMP and lysozyme in the *Ae. aegypti* midgut) were most strongly expressed in the proventriculi. This distribution of key genes leads us to hypothesize that the immune regionalization of the *An. gambiae* midgut is likely similar to that of *Ae. aegypti*. It should be noted, however, that the short amidase, *PGRPS3*, a putative negative regulator of the IMD pathway, is robustly expressed in the *An. gambiae* proventriculus, in striking contrast to its *Ae. aegypti* counterparts. We further note that, while the amplitude of *GAM1* and *LYSC7B*/*LYSC7* investment is comparable in the two species, the *An. gambaie* proventriculus and anterior midgut transcriptomes contained large quantities of transcripts for cecropins and defensins, rendering their overall AMP investment more than five times higher than the same regions in *Ae. aegypti* (S6B Fig). In both species, the posterior midgut’s investment in AMPs was negligible. Altogether, the expression of digestive (Fig 6B) and defensive genes (Fig 6C) as well as a GOEA (S6C Fig) of the *An. gambiae* midgut regions confirm that the midgut structure-function relationship is well conserved between the two mosquito species. Figure 6D provides a summary of enriched functions and prominently expressed gene families that are shared by the midgut regions of *Ae. aegypti* and *An. gambiae*.

### A small number of disparately expressed genes differentiate *Aedes aegypti* and *Anopheles gambiae* midgut transcriptomes

The conservation of gut functional regionalization may be due either to conserved expression patterns among orthologs across evolutionary distance, or to convergent patterning of non-homologous genes of shared function. To evaluate how well the relative expression of individual orthologous genes has been maintained between *Ae. aegypti* and *An. gambiae* midgut regions, we performed a series of correlation analyses using only the 7430 genes classified by Orthogroups Analysis (S8 Table) as one-to-one orthologs. Scaled expression values for the orthologs were poorly correlated both in whole guts (Fig 7A), and in dissected regions (S7.1A Fig), with R^2^ values ranging from 0.08 (proventriculus) to 0.56 (posterior midgut). However, the distribution of data points in the correlation graphs suggested that a few highly/disparately expressed genes substantially depressed correlation. To test the effect of these genes on overall correlation, we sequentially censored genes that were strongly and significantly disparately expressed between both species (DESeq2, padj<0.05), rescaled the expression of the remaining one-to-one orthologs, and plotted the resulting changes in slope and correlation coefficient (Fig 7B, S7.1B). For the proventriculus, anterior midgut, posterior midgut, and whole gut the censorship of, respectively 4, 10, 20, and 35 1-to-1 orthologous pairs was sufficient to reveal strong correlation between the remaining orthologs, with slopes ranging from 0.78 to 1.1 and R^2^ values ranging from 0.77 to 0.89. From this, we conclude that a ‘species signature’ masking the transcriptional similarity of *Ae. aegypti* and *An. gambiae* midguts is the product of a small number of highly expressed/disparately expressed genes.

**Fig 7.**
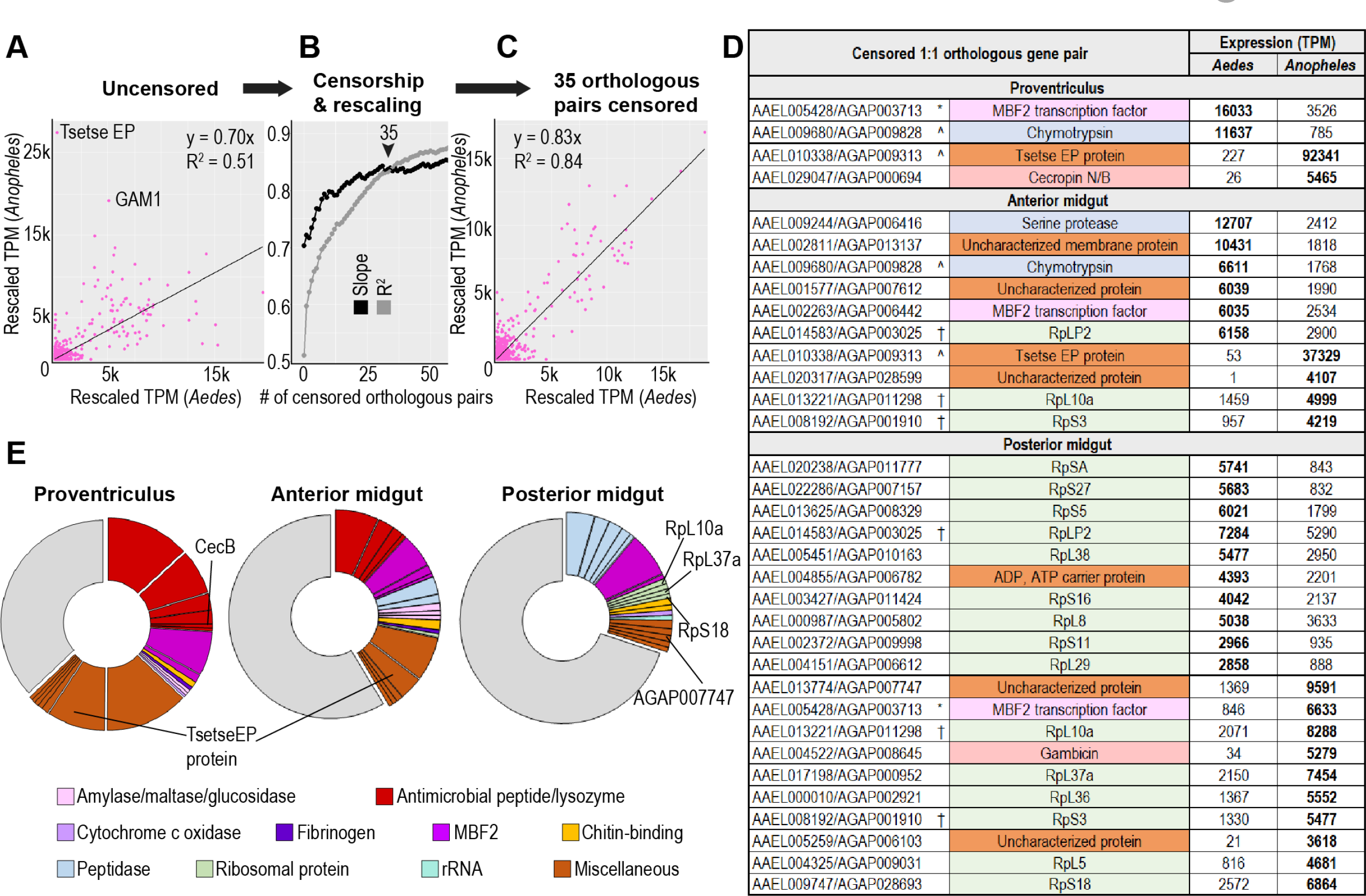
A small number of disparately expressed genes constitute a ‘species signature’ differentiating midgut transcriptomes in *Aedes aegypti* and *Anopheles gambiae*. (A-C) Correlation of one-to-one orthologous genes before and after censorship of 35 most disparately expressed orthologous pairs. (D) One-to-one orthologs censored in a regional correlation analysis; (^) signifies orthologous pairs censored in both proventriculus and anterior midgut, (†) signifies orthologous pairs censored in both anterior midgut and posterior midgut (*) signifies orthologous pairs censored in both proventriculus and posterior midgut. (E) 20 highest expressed transcripts per region, grouped by category/function.

Figure 7D displays the genes censored from our correlation analysis in the three midgut regions. The most disparately expressed gene in both the *An. gambiae* proventriculus and anterior midgut encodes an uncharacterized protein containing a domain (IPR007931) from the Tsetse EP protein which, in tsetse fly species has been shown to play a role in resisting the establishment of trypanosome infections [65]. MBF2 transcription factors were also censored in each region: twice they were expressed higher in *Ae. aegypti* (proventriculus and anterior midgut) and once in *An. gambiae* (posterior midgut). Among the disparately expressed orthologs, the most abundant category was ribosomal proteins. We observed that in both the anterior and posterior midgut, *Ae. aegypti* had significantly increased the expression of some ribosomal proteins relative to their one-to-one orthologs in *An. gambiae*, and *vice versa*. Phylogenetic analysis of ribosomal proteins (S7.2A Fig) confirmed that the putatively one-to-one orthologous ribosomal genes identified by Orthogroups Analysis were sister sequences and that the disparately expressed pairs of orthologous genes were widely distributed throughout the phylogenetic tree. We therefore conclude that evolutionary changes between the two species have reshuffled the expression of ribosomal proteins, significantly reducing the roles of some while increasing those of others.

Many of the genes censored in our correlation analysis were among the highest expressed genes in the three *Ae. aegypti* and *An. gambiae* midgut regions. A follow-up comparison of the twenty highest expressed genes in the midgut regions of the two species (Fig 2C, Fig 7E) revealed striking differences in both amplitude and function. In the proventriculus, *An. gambiae* devoted a higher proportion of its transcriptome to the expression of a small number of genes. The three highest expressed, including an uncharacterized gene (AGAP013543), the Tsetse EP gene (AGAP009313), and a cecropin (*CEC2*) together composed more than one third of the transcriptome. Both species’ proventriculi shared expression of two MBF2 transcription factors (AAEL005428 and AAEL013885 in *Ae. aegypti*, AGAP011630 and AGAP000570 in *An. gambiae*), as well as the AMP gambicin (*GAM1*). However, the *An. gambiae* proventriculus also expressed three cecropins (*CEC2*, *CEC1*, *CECB)*, one defensin (*DEF1*) and a lysozyme (*LYSC7*) among its top twenty genes, together totaling more than 25% of its transcriptome. Cecropins (*CEC1*, *CEC2*) and a defensin (*DEF1*) were likewise among the twenty highest expressed genes in the *An. gambiae* but not the *Ae. aegypti* anterior midgut. In the posterior midgut, the highest expressed peptidases in *An. gambiae* were much lower expressed than their counterparts in *Ae. aegypti*, and the top twenty genes in the *An. gambiae* posterior midgut included two MBF2 transcription factors (AGAP000570, AGAP003713) that were absent among the highest expressed genes in the corresponding region in *Ae. aegypti.* Altogether, we find that while gut regional specializations have been broadly conserved between *Ae. aegypti* and *An. gambiae*, and while most one-to-one orthologous genes maintain closely correlated expression patterns, this correlation is lost among a few of the highest expressed genes in the midgut, creating a ‘species signature’ that differentiates the two midguts’ transcriptomes.

## Discussion

Aegypti-Atlas is a new resource which contextualizes and dissects the transcriptome of the sugar and blood-fed *Aedes aegypti* gut. Aegypti-Atlas will allow exploration of gene expression across the main parts/regions of the female mosquito body/gut in an easily accessible online database. In addition, we have performed a first analysis of this new dataset, examining how biological functions are distributed in the parts of the mosquito body and the regions of the gut, as well as how they change in the gut over the course of blood meal digestion. Our study demonstrates the importance of estimating investment, yield, and output in transcriptomic studies. In addition, this research has yielded new insights into such diverse areas as the regionalization of digestive function, the temporal patterning of peptidase transcription, the organization of immune defenses, and the similarities and differences between the guts of two hematophagous vectors.

### Assessments for Highest Expressed Genes and Categorical Investment/Output are Valuable Complements to GO Enrichment Analysis

In bioinformatic analysis of large-scale transcriptomic data sets, GOEA is often the preferred method for determining the functional role of a tissue or body part, or for detecting functional differences across conditions. We have employed GOEA to derive valuable information about the enrichment of functions in mosquito body parts, gut regions, and blood-fed conditions. However, enrichment analysis alone cannot capture all the complexities of comparative transcriptomics, as it is primarily sensitive not to quantities of transcripts from a given category, but to quantities of genes belonging to that category. Furthermore, GOEA is substantially less robust in non-model species, owing to incomplete annotations which impact many key genes. We propose that complementary approaches, such as identifying highest expressed genes and examining transcriptomic profiles through the lenses of *investment* and *output*, can help to strengthen conclusions and capture dynamics which might otherwise be overlooked. By examining the twenty highest expressed genes in each body part and gut region, we were able to identify genes that account for large proportions of the transcriptome, but which were missed by enrichment analysis (*e.g.*, *GAM1* in the proventriculus, MBF2 factors in the anterior midgut). Comparing the top-expressed genes across transcriptomes allowed us to see that some body parts are dominated by a small number of highly expressed genes (*e.g.*, the gut) while others spread their transcriptome more equally over the genome (*e.g.*, the ovaries). Comparing the relative expression of one-to-one orthologs between species allowed us to identify genes that are disproportionately expressed in one species (*e.g.*, the Tsetse EP protein, a putative antimicrobial effector in the *An. gambiae* proventriculus) and to detect a reshuffling of expression among ribosomal proteins across the two species.

Another approach we adopted – summing transcripts belonging to a functional category and comparing the sums across body parts, regions, or timepoints – allowed us to assess relative investment in the given category. We confined the use of this method to categories of genes encoding proteins that are all presumed to have similar molecular function, narrowly defined (*e.g.*, peptidases cleaving proteins, or antimicrobial peptides killing pathogens). The utility of evaluating regional or temporal specialization by summing transcripts (investment) as opposed to counting genes (enrichment), is evident when we consider that using GOEA, the highest expressed peptidase in the gut carries no more weight than the fifth highest – despite a difference of over 165,000 TPM, or 16.5% of the total gut transcriptome. Using this investment approach, we were also able to break large categories (*e.g.*, peptidases) into smaller categories (*e.g.*, carboxypeptidases, dipeptidases, *etc.*) for a fine-grained picture of where and when each was most expressed. There is, however, an important caveat to the interpretation of cumulative analysis as it fails to account for differential transcriptional yields between profiled body parts, regions, or conditions. We overcame this caveat by weighting transcriptional investment by RNA yield to calculate estimated regional/temporal output of digestive transcripts, from which we were able to infer how certain tasks (*e.g.*, sugar digestion) are divided between the regions of the gut, as well as how peptidase transcript output changes over the course of blood meal digestion.

### Spatial organization of gut function is conserved across mosquito species; few highly/disparately expressed genes drive transcriptomic differences

The *Ae. aegypti* gut is linearly divided into five anatomically distinct regions. With the caveat that the crop only admits sugar meals, ingested materials encounter each of these regions sequentially. The regions of the gut therefore possess an ordinal quality, corresponding to stages (either transient or prolonged) of the ingestion/digestion process. Through GOEA and quantitative evaluation of transcripts belonging to digestive categories, we discerned a clear pattern of strong investment in sugar digestion/absorption in the anterior midgut, as hypothesized by Hecker [66]. Peptidases, by contrast, were predominantly expressed in the posterior midgut. These specializations were maintained under blood-fed conditions, and largely conserved in *An. gambiae* midguts. Other notable instances of conserved regional function are apparent in the proventriculus, where both species share enrichment of wingless signaling components [31] and AMPS [67] with other dipteran species.

While the midgut regions of *Ae. aegypti* and *An. gambiae* shared many important functions, and their one-to-one orthologs maintained strikingly close transcriptional correlation, we noted that differences in the expression of a small number of highly expressed genes created large disparities in their overall categorical apportionment of transcripts. Most notably, the *An. gambiae* proventriculus and anterior midgut expressed far greater quantities of AMPs – possibly in response to some microbial presence - as well as a handful of highly expressed but poorly characterized genes, including an apparent ortholog of a tsetse midgut protein with a defensive function [65]. Meanwhile, the *An. gambiae* posterior midgut expressed far fewer peptidase transcripts than its *Ae. aegypti* counterpart, and far more of an MBF2 family of transcription cofactors which, in the *Ae. aegypti* gut, was only prominent in the anterior regions of the sugar-fed midgut and, intriguingly, the blood-fed posterior midgut. In Drosophila and *Bombyx mori*, MBF2 factors have been shown to complex with the transcription factor FTZ-F1 and cofactor MBF1 to activate transcription of target genes [68, 69]. We cannot say whether this role is conserved in *Ae. aegypti* and *An. gambiae*. However, the expression of MBF2 factors in both species outstripped the expression of their orthologs of *FTZ-F1* (AAEL019863, AAEL026810, AGAP005661) and *MBF1* (AAEL008768, AGAP004990) by several orders of magnitude in multiple conditions and/or gut regions (see Aegypti-Atlas website), suggesting MBF2 may not be confined to cofactoring FTZ-F1 in these species. Altogether, this analysis yielded multiple ‘species signature’ genes worthy of closer scrutiny and functional study.

### The anterior midgut participates in the transcriptional response to blood feeding

The anterior midgut has been held to play little or no role in the process of blood meal digestion, as the blood bolus is sealed into the posterior midgut by the formation of the peritrophic matrix shortly after ingestion [70]. However, late in the blood meal response (24 hrs) we found that the anterior midgut had increased its investment in some categories of digestive enzymes and transporters (*e.g.*, sphingomyelinases, sterol transporters, amylases/maltases, glucosidases, Fig 4D) as well as its overall transcriptional yield (Fig 4E). While the proportional output of the anterior midgut drops relative to the posterior midgut (Fig 4F), it still contributes approximately one third of the transcripts for sugar-digesting enzymes in the whole midgut (exclusive of proventriculus). This finding is congruent with Billingsley and Hecker’s observation that alpha-glucosidase activity was elevated in homogenates of *Anopheles stephensi* anterior midguts at 24 hrs pbm [71]. As the majority of amylases/maltases and glucosidases in the *Ae. aegypti* genome possess signal peptides but lack transmembrane domains, it is possible that enzymes secreted in the anterior midgut are capable of diffusing into the extra-peritrophic area in the posterior midgut where they may participate in blood meal digestion.

### Gut peptidases are dynamically up and downregulated in sequential transcriptional waves upon blood feeding

Peptidase expression in the *Ae. aegypti* gut unfolds in a series of phases, coordinated by rising and falling titers of JH and ecdysone. In the post-emergence/pre-vitellogenic phase, JH drives the transcription of a cohort of “early” peptidases, priming the gut for its first blood meal [39,40,42,72]. Within hours of blood feeding, translation of this pool of transcripts is initiated and peptidase activity begins to increase [37]. Next, rising ecdysone titers drive the transcription of a “late” cohort of peptidases, which complete the digestive process [48]. Finally, early peptidases are transcribed anew [38, 40] amid a postprandial surge of JH [73–76], restoring the gut to a state of readiness for its next blood meal.

Peptidase activity peaks somewhere between 18 and 36 hours pbm [43,44,47], as do transcripts for some of the highest expressed “late” phase peptidases [49,50,52]. However, a few publications have documented peptidases peaking at earlier timepoints [51–53], and that at least one of these (*SPI*) is directly responsive to/dependent on ecdysone signaling [48]. Here, our genome-wide multi-timepoint series has demonstrated that these early-peaking genes are not isolated and exceptional but are, rather, members of large transcriptional cohorts. We show that peptidases in the blood-fed *Ae. aegypti* gut are expressed not only in two phases: “early” (translational) and “late” (transcriptional), but that the “late” transcriptional response manifests in multiple successive waves or shifts. We have divided these into three (“rapid”, “intermediate”, and “delayed”), but because we only obtained RNAseq profiles for guts at baseline and three blood-fed timepoints, we can only speculate how many of these shifts are actually triggered over the course of blood meal digestion, and how synchronously any of the peptidases that our clustering analysis grouped together are actually regulated. Our timepoints were too broadly spaced to say at what time each gene reached its true peak, or whether peptidases that were apparently quiescent throughout blood meal digestion in fact participated in a shift that was too transient to be detected by our experimental design. Future work will likely uncover even more complexity than we have described.

The sharp temporal patterning of peptidase transcription in the blood-fed mosquito gut raises intriguing questions. How is this transcriptional choreography achieved? Is the transcription of all “late” phase peptidases ecdysone-dependent, and are rising ecdysone levels sufficient to drive them? To what extent is the transcriptional induction of succeeding waves dependent on the activity of prior ones? Why are some “delayed” peptidases so slow to show any transcriptional response when ecdysone titers are ascendent (*e.g.*, AAEL008782, Fig 3E)?. And what mechanism effects the precipitous drop in the “rapid” cohort of induced peptidases (ascendent at 6 hrs pbm) at a timepoint (24 hrs pbm) when ecdysone titers are still high (*e.g.*, AAEL013715) [77]? What is the adaptive significance of the rapid up and down-regulation of so many genes of shared function? We observed that the gut’s investment in aminopeptidases, dipeptidases, and carboxypeptidases has distinct temporal peaks, suggesting that polypeptides may be attacked more from one terminus or the other at different times over the course of digestion. However, most of the peptidases that participate in shift-changes are endopeptidases, and it is not clear how the sequential substitution of one set of endopeptidases for another – as opposed to simultaneous expression - helps to move the process of digestion forward. We speculate that rapidly changing conditions in the blood-fed gut may alter the stability and kinetics of specific peptidases, and that different peptidases predominate at different times because they are adapted to function in specific, transient conditions obtaining at corresponding stages of the digestive process.

### The anterior and posterior midgut regions are, respectively, characterized by signs of immune activity and immune tolerance

In the midguts of both *Ae. aegypti* and *An. gambiae*, the majority of AMP transcripts are contributed by the proventriculus and anterior midgut (Fig 5D, S6B Fig) with minimal expression in the posterior midgut. Also, in both species, the transcription factor *caudal* which, in Drosophila [64] and *An. gambiae* [63] has been shown to repress AMP expression, is dominantly expressed in the posterior midgut. The expression of immune activating recognition proteins (*PGRPLC* and, in *Ae. aegypti*, *PGRP-LE*) is less profoundly patterned, but still displays some tropism toward the anterior midgut in both mosquitoes. Altogether, these patterns suggest that ingested microbes are subjected to heightened immune surveillance and antimicrobial activity in the first regions of the midgut but are thereafter well tolerated by the posterior midgut, where they may serve some mutualistic functions. We speculate that selection by gambicin in the proventriculus and anterior midgut of *Ae. aegypti* and *An. gambiae* may play an important role in determining the composition of the microbiota that seed the posterior midgut (S7.3A Fig).

### Holotricin and gambicin are highly expressed under baseline conditions in mutually exclusive regions of the body

Our examination of the expression of immune effectors yielded the observation that at baseline, AMP expression in *Ae. aegypti* is dominated by two genes: holotricin and gambicin. We observed that holotricin predominated in the carcass, while gambicin was highly expressed in the gut and Malpighian tubules. While it is not uncommon for antimicrobial effectors to display strong tropisms to specific tissues, we are not aware of comparable instances where a single AMP exhibits such overwhelming transcriptional dominance. In future work, it might be interesting to compare the activity of holotricin and gambicin against different types of microbes (Gram-positive, Gram-negative, fungi, *etc.*), and to examine the effects that silencing these genes has on midgut communities, and on mosquitoes’ survival in the contexts of oral and systemic infection.

## Conclusion

In this manuscript we introduce Aegypti-Atlas, a repository of RNAseq data, and demonstrate how these data can yield insights into tissue function, the organization of digestive specializations, regulatory networks, and immune effectors, as well as the changes wrought by diet and evolution in mosquitoes. This resource may also be useful for the creation of functional genomic tools (*e.g.*, tissue-specific expression systems) which will afford researchers a greater degree of control in experiments and allow for more confidence in the interpretation of results. It is our hope that Aegypti-Atlas will be valuable to other investigators in many areas of mosquito biology.

## Methods

### Mosquito provenance and rearing

For all experiments, we used 5 to 12 day-old mated female mosquitoes of the Thai strain (*Ae. aegypti*) or the G3 strain (*An. gambiae*) kindly provided by Laura Harrington. Mosquitoes were reared at a density of 200 larvae per 1-liter tray. *Ae. aegypti* received 720 mg of fish food (Hikari #04428). *An. gambiae* received 50 mg of ground fish food per day from day 1-4 of development, and 150 mg per day thereafter until pupation. Adults were maintained in humidified chambers at 29°C on a diet of 10% sucrose *ad libitum*.

### Blood feeding and mock blood feeding

For all RNA-seq experiments involving blood-fed guts, mosquitoes were starved for twenty-four hours, then blood-fed on live chickens without anesthetic in accordance with Cornell University IACUC approved protocol #01-56. For our extended blood feeding time series by RT-qPCR, mosquitoes were fed through a membrane on rooster blood treated with the anticoagulant sodium citrate (Lampire Biological Laboratories, Pipersville, PA, catalogue #7208806). For experiments where RNA yield was estimated in guts and midgut regions under blood-fed conditions, we employed an artificial formulation (SkitoSnack) [78] fed through a membrane. This formulation was used in order to avoid any skewing of yield which might result from RNA content in a natural blood meal. For RNA yield experiments, boluses were removed during dissection.

### Dissections & RNA extraction

Mosquitoes were sacrificed by submersion in 70% ethanol and dissected in sterile PBS. For body part samples, the last 2-3 abdominal segments were removed to eliminate the sperm-containing spermatheca. Whole guts, Malpighian tubules, and ovaries were then dissected out. The residual carcass was divided into head, thorax, and abdomen. A small anterior section of the thoracic carcass containing the salivary glands was excised to exclude salivary gland transcripts from thoracic profiles. For RNAseq experiments, a minimum of 20 individuals per replicate was used for large body parts (*e.g.*, whole body, abdominal carcass, whole guts) and up to 500 individuals per replicate for small parts (*e.g.*, crop, proventriculus). For RT-qPCR time-series experiments, a minimum of 10 individuals was dissected per replicate. For all experiments, a minimum of 3 replicates was prepared per part and/or condition. All samples were homogenized in TRIzol and stored at -80°C. For RNAseq experiments, RNA was extracted via a modified phenol-chloroform method [79]. For RT-qPCR time series and RNA yield experiments, a standard phenol-chloroform extraction was performed, and RNA concentration was evaluated by Qubit fluorometer.

### Library preparation & sequencing

Libraries were prepared using the Quantseq 3’ mRNA-seq prep kit from Lexogen according to the manufacturer’s instructions. Sample quality was evaluated before and after library preparation using a fragment analyzer (Advanced Analytical). Libraries were pooled and sequenced on the Illumina Nextseq 500 platform using standard protocols for 75bp single-end read sequencing at the Cornell Life Sciences Sequencing Core. Sequences have been deposited on NCBI (BioProject ID: PRJNA789580).

### Analysis pipelines

Quality control of raw reads was performed with fastqc (https://github.com/s-andrews/FastQC) and reads were trimmed by BBMap (https://jgi.doe.gov/data-and-tools/bbtools/) and then mapped to the *Ae. aegypti* transcriptome (version l52) or the *An. gambiae* transcriptome (version P4.12) using Salmon version 0.9.1 with default parameters. Salmon’s transcript-level quantifications were then aggregated to the gene level for gene-level differential expression analysis with the “tximport” package on the R version 3.5. DEseq2 was used for differential expression analysis and PCA analysis. GOEA was performed using the TopGO package.

All phylogenetic trees were constructed using Geneious software version R11, with the following settings. Alignment type: Global alignment, Cost Matrix: Blosum62, Genetic Distance Model: Jukes-Cantor, Tree Build Method: Neighbor-Joining. Clustering was performed using the Pheatmap package in R with gene expression scaled by row. Z-scores were calculated by first censoring all genes expressed at less than 2 TPM in the relevant body parts or gut regions, then employing the following expression for each gene in each body part or gut region: z = (x - µ)/s, where x is the expression in the body part or gut region, µ represents the mean expression of all body parts or gut regions, and s is the standard deviation. The peptide sequences of AaegL5.2 and AgamP4.12 were used to find orthogroups and orthologs in the two species using OrthoFinder v2.3 [80]. The longest transcript variant of each gene was extracted to run the OrthoFinder with the following options: “-M msa -S blast -A muscle -T iqtree”.

### Calculating the “mosquito equation”

If X is defined as the expression of a specific gene in a given body part (in TPM), and the part’s initial stands for the relevant scaling factor, the relationship between the expression of that gene in each body part versus in the whole body can be expressed as:

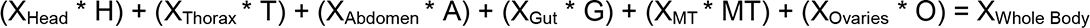

Where H + T + A + G + MT + O = 100% Whole Body expression For marker genes the expression in other body parts may be considered to be negligible, (*i.e.*, for a head marker, X_Thorax_, X_Abdomen_, X_Gut_, X_MT_, and X_Ovaries_ are all ≈ 0). Therefore:

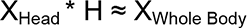

We solved for the estimated scaling factor for each of our qualifying markers, then averaged the values we obtained for each body part to create a rough estimate of the percent of transcripts in the whole body that are contributed by each part.

### RT-qPCR

For all RT-qPCR amplifications, RNA was pretreated with DNase (Quantabio 95150-01K) and cDNA was prepared using the qScript cDNA synthesis kit (Quantabio 95047-100) and amplified using Perfecta SYBR Green Fastmix (Quantabio 95072-012) in a Bio-Rad CFX-Connect Instrument. The primer sequences used in this study are available in S9 Table.

## Acknowledgements

Special thanks to Jonathan Revah for creating the Aegypti-Atlas website, and to Laura Harrington, Sylvie Pitcher, and Garrett League for their kind provision of Thai and G3 strain mosquitoes. We also acknowledge members of the Buchon lab, Kristin Michel, Courtney Murdock, Laura Harrington, and Jeffrey Scott for helpful comments on the manuscript.

## Supplemental Figure Legends

**S1.1 Fig.**
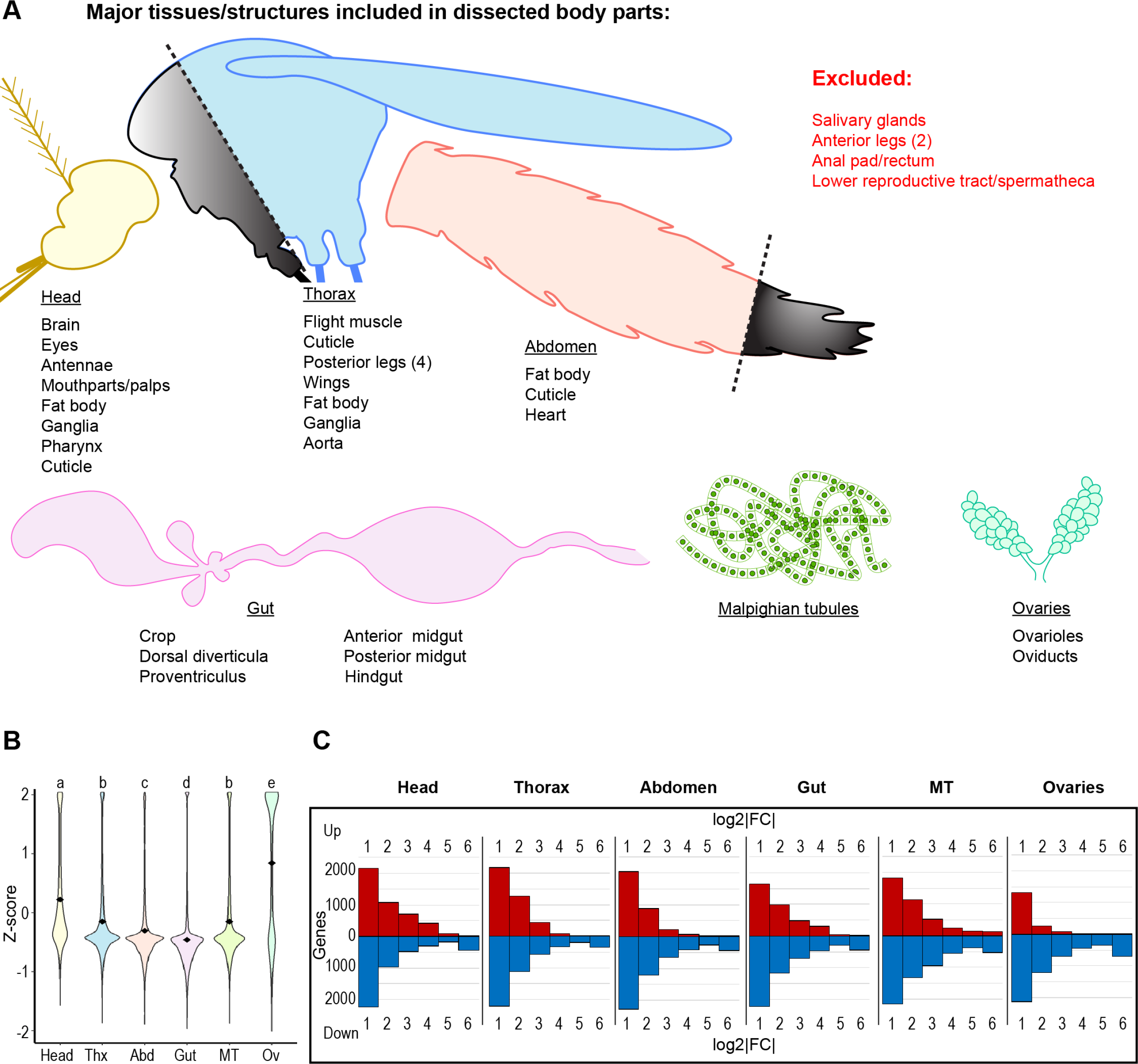
Ovaries possess many highly enriched genes relative to other body parts, but few relative to whole body. (A) Diagram of included/excluded structures/tissues in dissected body parts (B) Transcriptome-wide z-scores for gene expression by body part. Data are z-scores for genes expressed ≥ 2 TPM in one or more body parts; black diamonds mark mean z-scores; statistics: One-way ANOVA, Tukey HSD. (B) Transcriptomic variation from whole body expression by body part (fold change adjusted by DESeq2, no significance criterion applied).

**S1.2 Fig.**
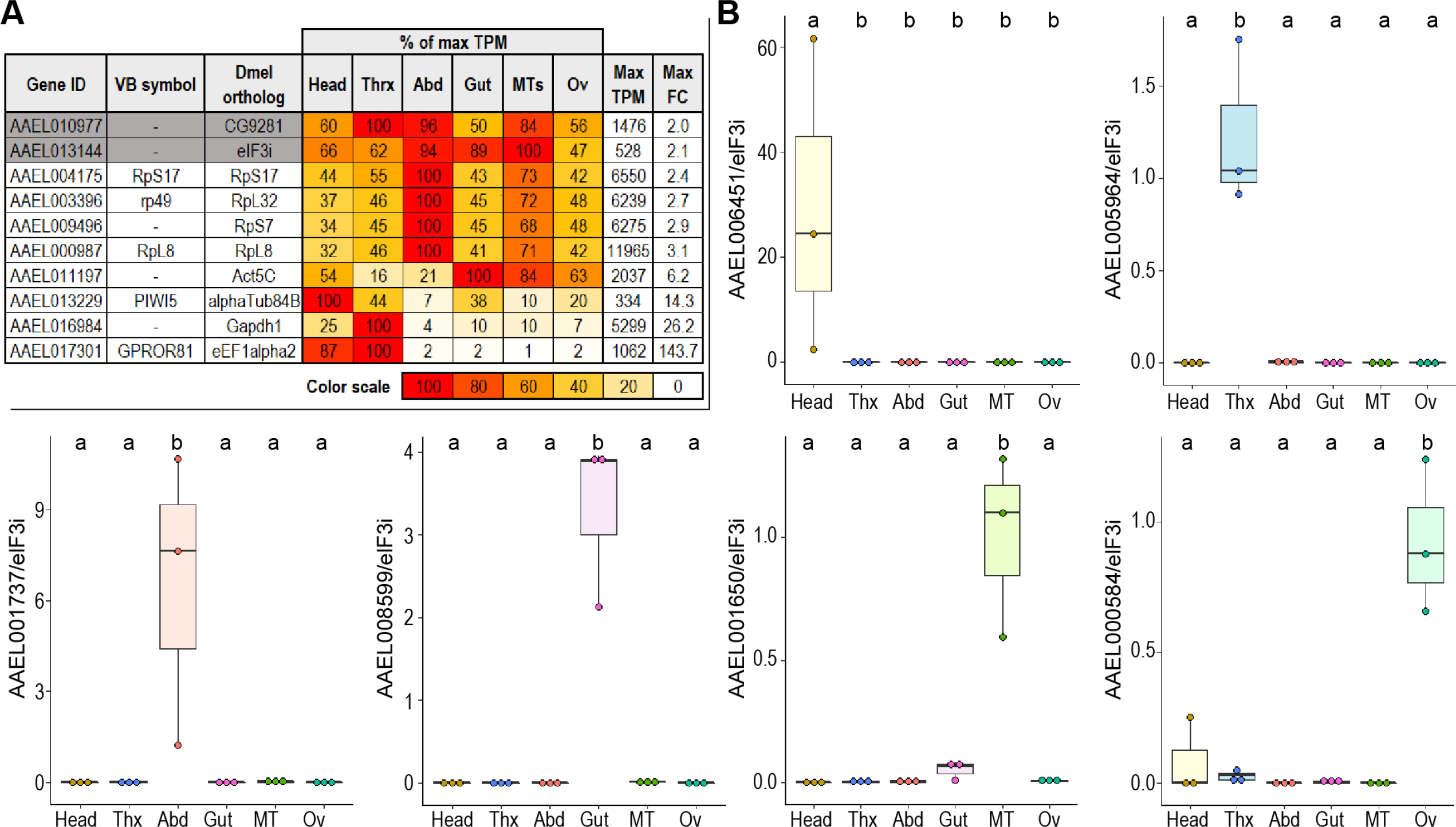
RT-qPCR confirms the anatomical specificity of putative marker genes. (A) Comparison of candidate reference genes for RT-qPCR analysis of putative marker genes; expression values are scaled as a percentage of the maximum expression observed (in TPM). (B) RT-qPCR validation of putative marker gene expression; reference gene: AAEL013144; 3 replicates; statistics: One-way ANOVA, Tukey HSD.

**S1.3 Fig.**
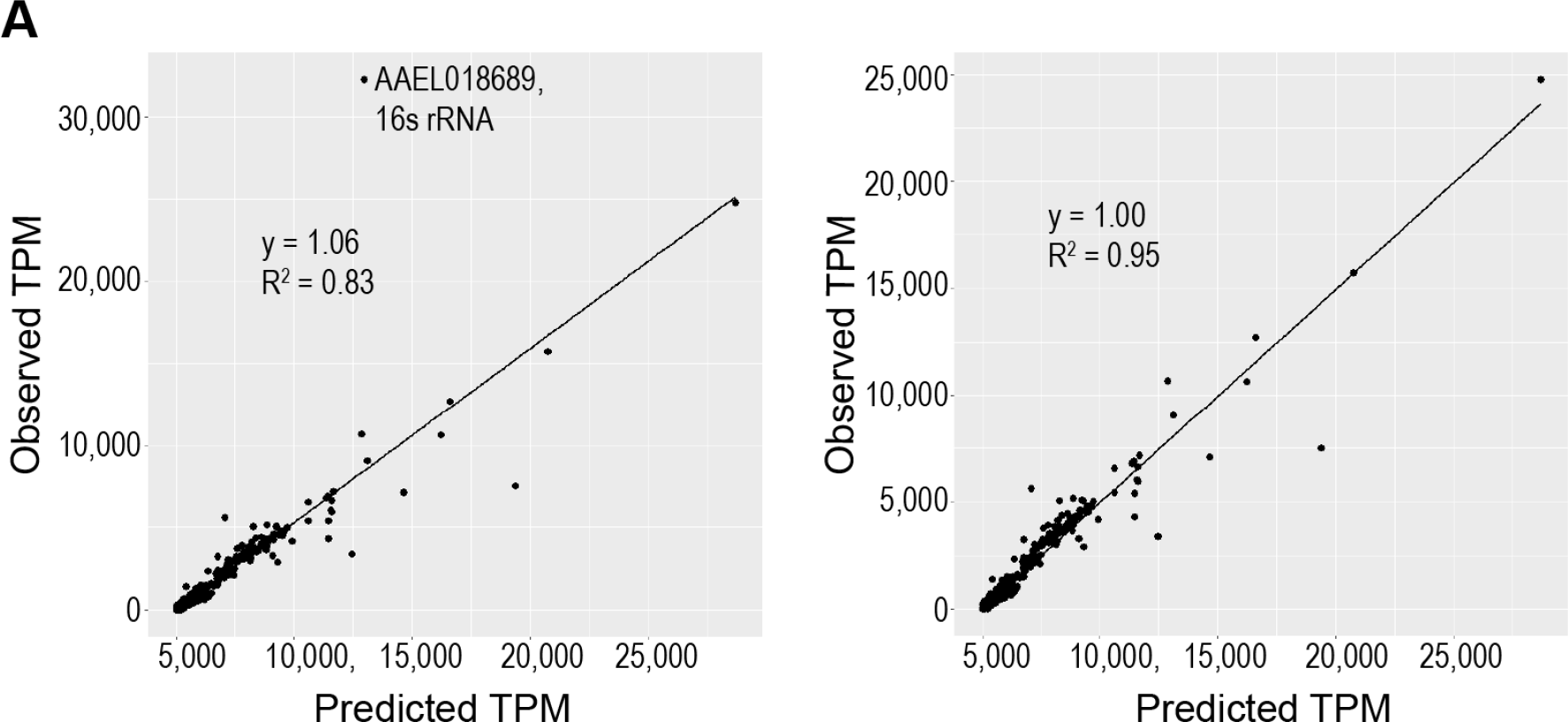
Calculated scaling factors accurately predict whole body gene expression. (A) Validation of estimated scaling factors for body part transcript contribution to whole body expression, with (left) and without (right) outlying gene.

**S2.1 Fig.**
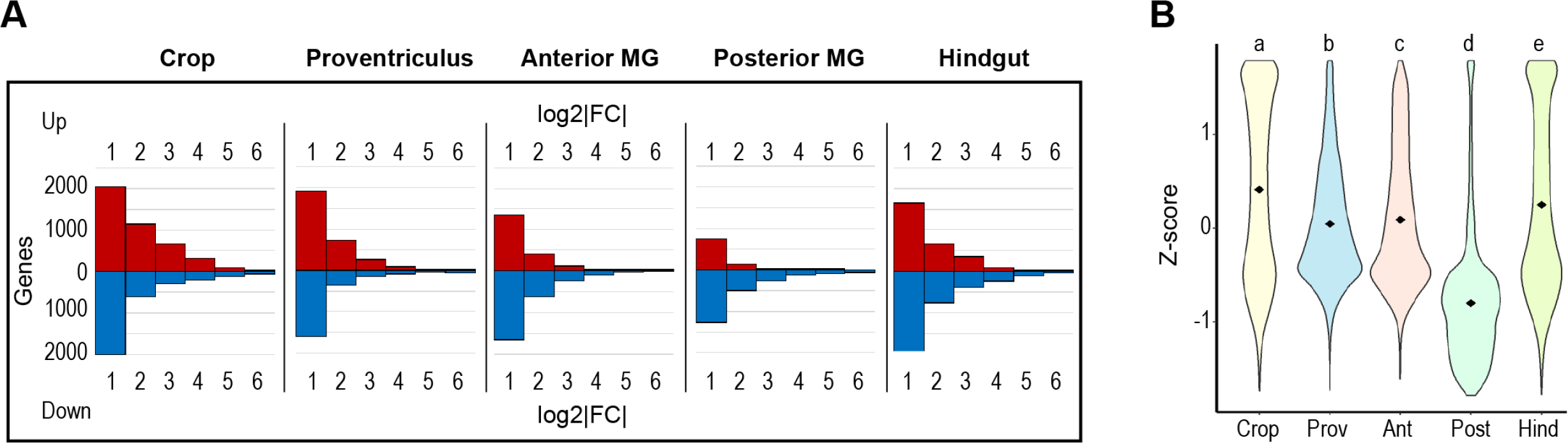
The transcriptome of the posterior midgut displays little deviation from the whole-gut transcriptome and low transcriptomic diversity. (A) Transcriptomic variation from whole gut expression by gut region (fold change adjusted by DESeq2, no significance criterion applied). (B) Transcriptome-wide z-scores for gene expression by gut region. Data are z-scores for genes expressed ≥ 2 TPM in one or more regions; black diamonds mark mean z-scores; statistics: One-way ANOVA, Tukey HSD.

**S3.1 Fig.**
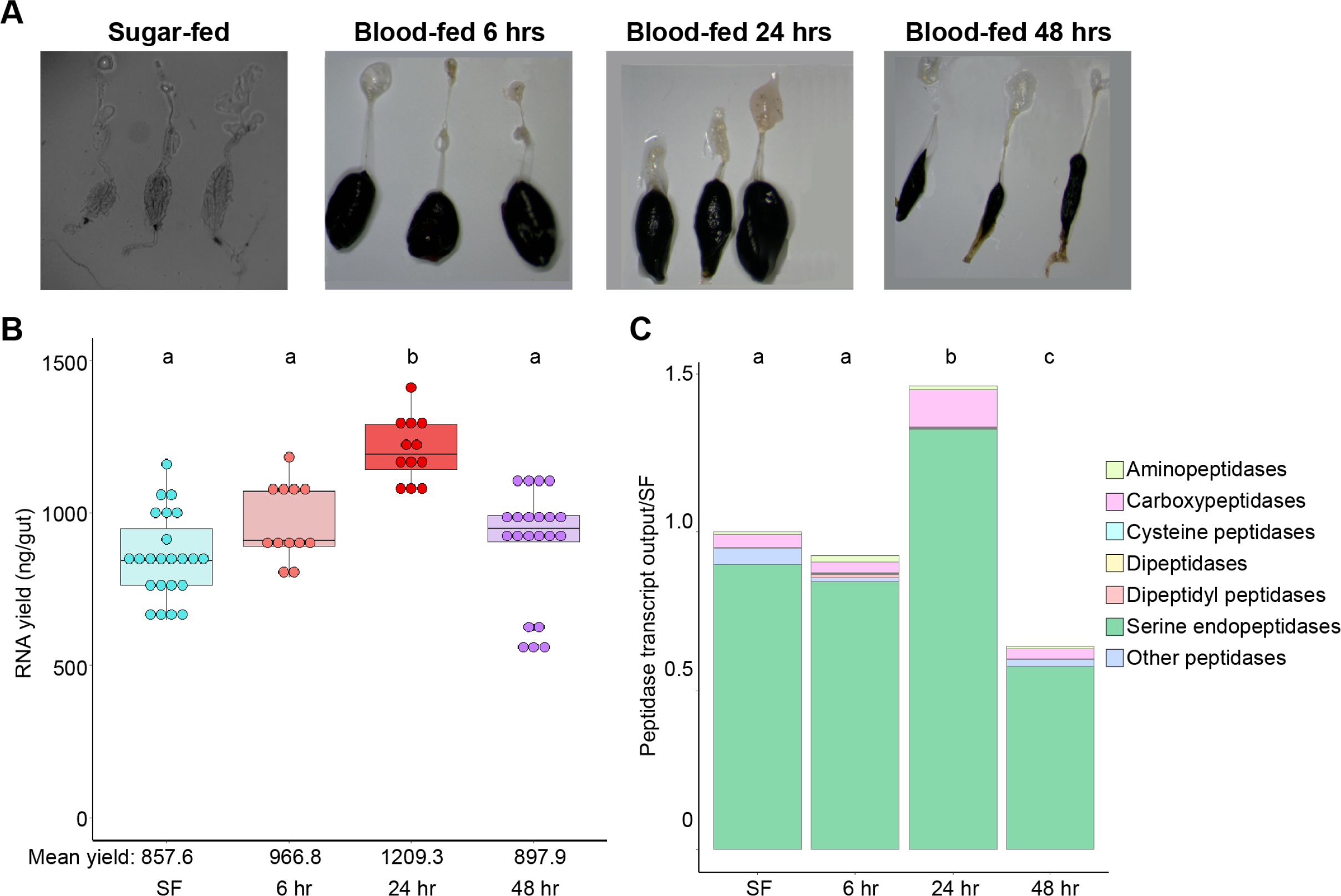
Increasing RNA yield drives higher gut peptidase output at 24 hours post blood meal. (A) Dissected guts from sugar-fed mosquitoes and mosquitoes at 6, 24, and 48 hrs pbm. (B) Estimates of gut’s transcriptional output in sugar-fed mosquitoes and at 6, 24, and 48 hrs pbm by empirical quantification of RNA yield; statistics: One-way ANOVA, Tukey HSD. (C) Estimated output of peptidases by category (categorical investments weighted by RNA yield, scaled to SF output); statistics: One-way ANOVA, Tukey HSD.

**S3.2 Fig.**
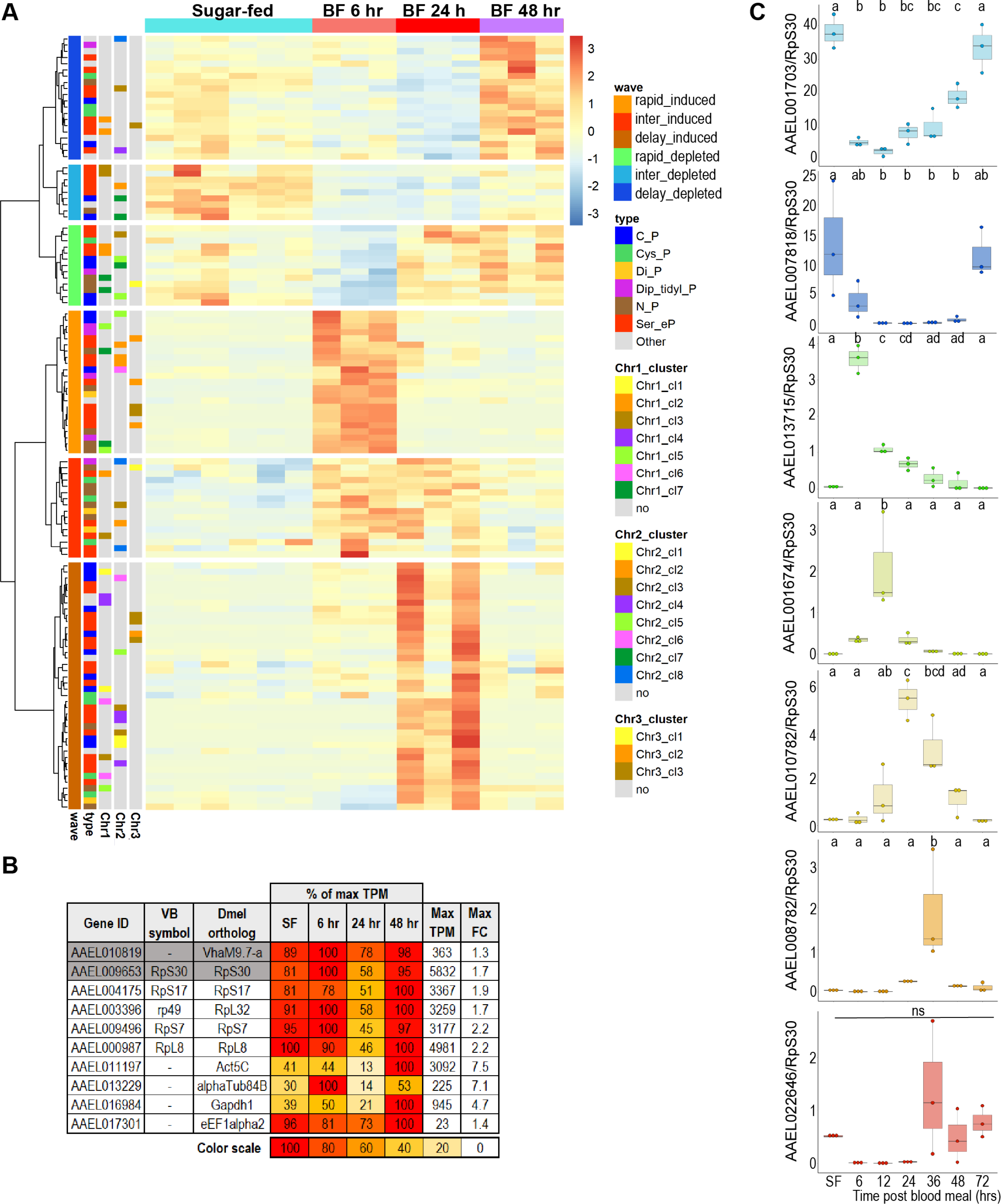
Gut peptidase transcripts are induced and depleted asynchronously during blood meal digestion. (A) Clustering analysis of the expression of peptidases across all timepoints of blood feeding time series. (B) Comparison of candidate reference genes for RT-qPCR analysis of peptidases in blood-fed guts; expression values are scaled as a percentage of the maximum expression observed (in TPM). (C) Unscaled results of RT-qPCR for selected peptidases relative to reference gene (AAEL009653); 3 replicates; statistics: One-way ANOVA, Tukey HSD.

**S3.3 Fig.**
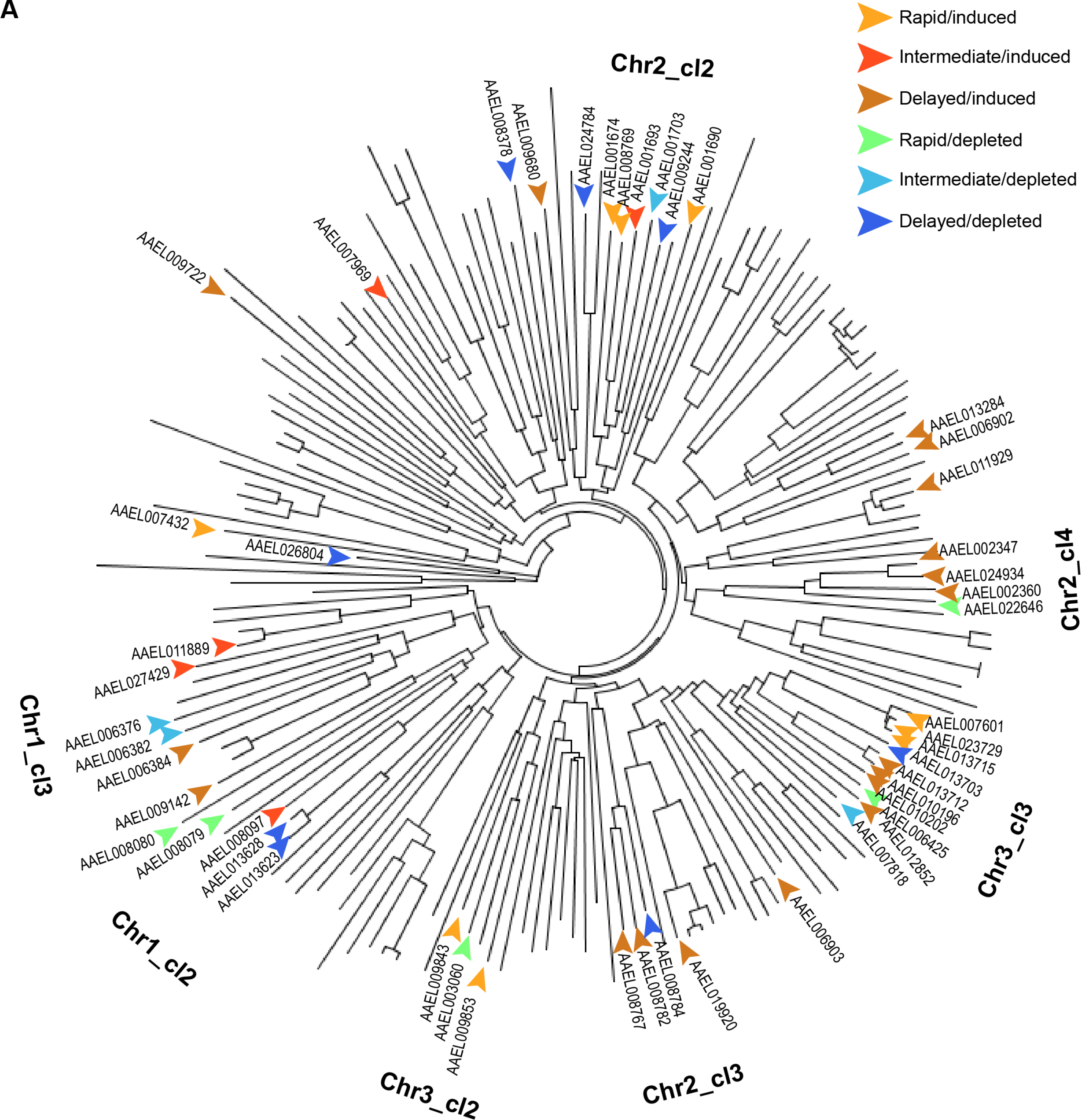
Temporal clustering of serine endopeptidases is independent of physical proximity and sequence similarity. (A) Phylogenetic tree for serine endopeptidases in *Aedes aegypti* with regulated genes labeled by temporal cluster; Note: superimposed chromosomal cluster labels indicate where a plurality of genes from that locus are located on the diagram, but not all genes proximal to a label belong to that chromosomal locus

**S6 Fig.**
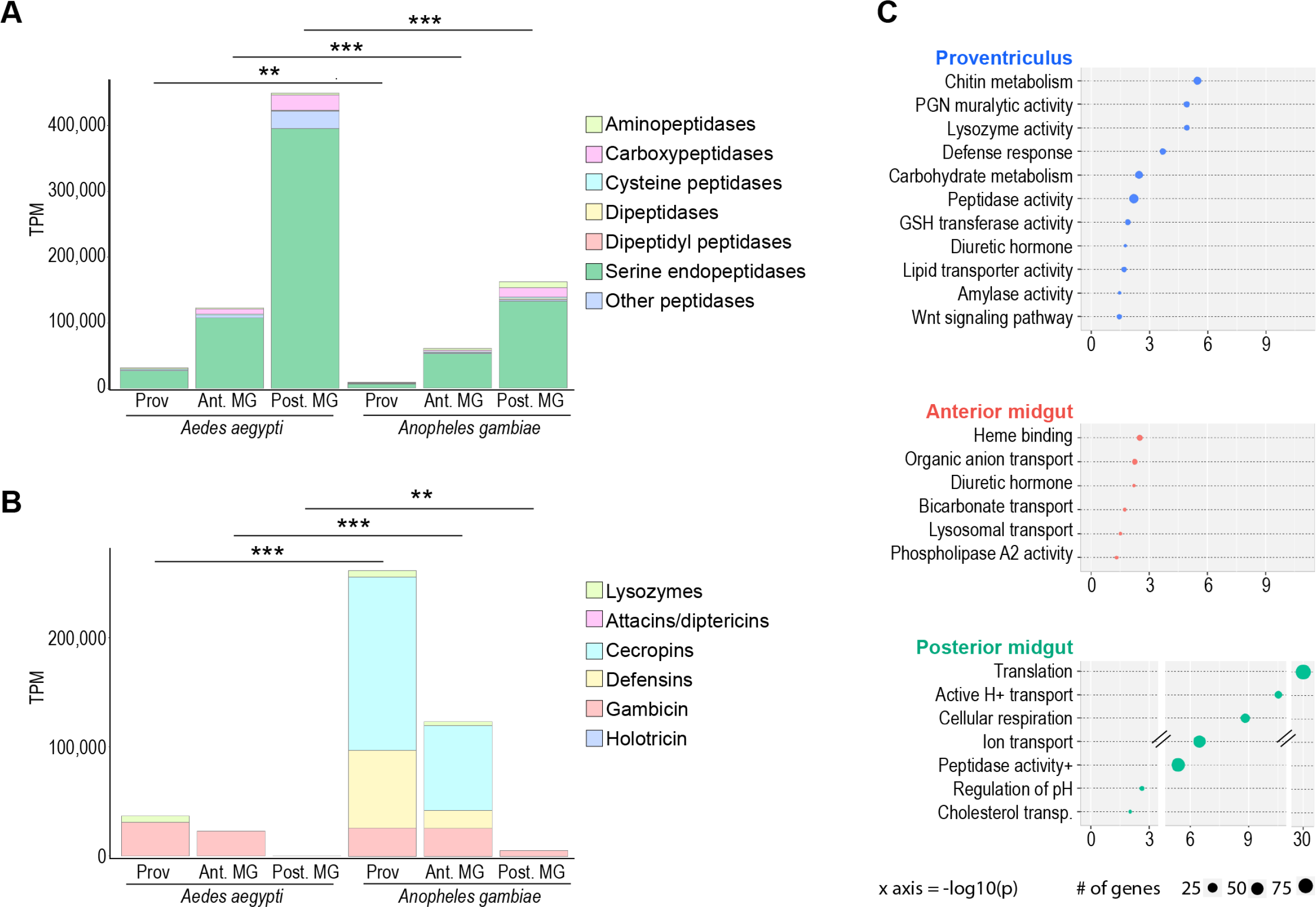
Midgut regions in *Aedes aegypti* and *Anopheles gambiae* display similar regional specializations, but disparate amplitudes of investment in key functions. Midgut regional investments in (A) digestive peptidases and (B) AMPs and lysozymes in *Ae. aegypti* and *An. gambiae*; statistics: unpaired t-test. (C) TopGO GOEA of regionally enriched genes (5x over whole gut for proventriculus, and anterior MG, 5x over other midgut regions for posterior MG) in *Anopheles gambiae* (DESeq2, padj <0.05).

**S7.1 Fig.**
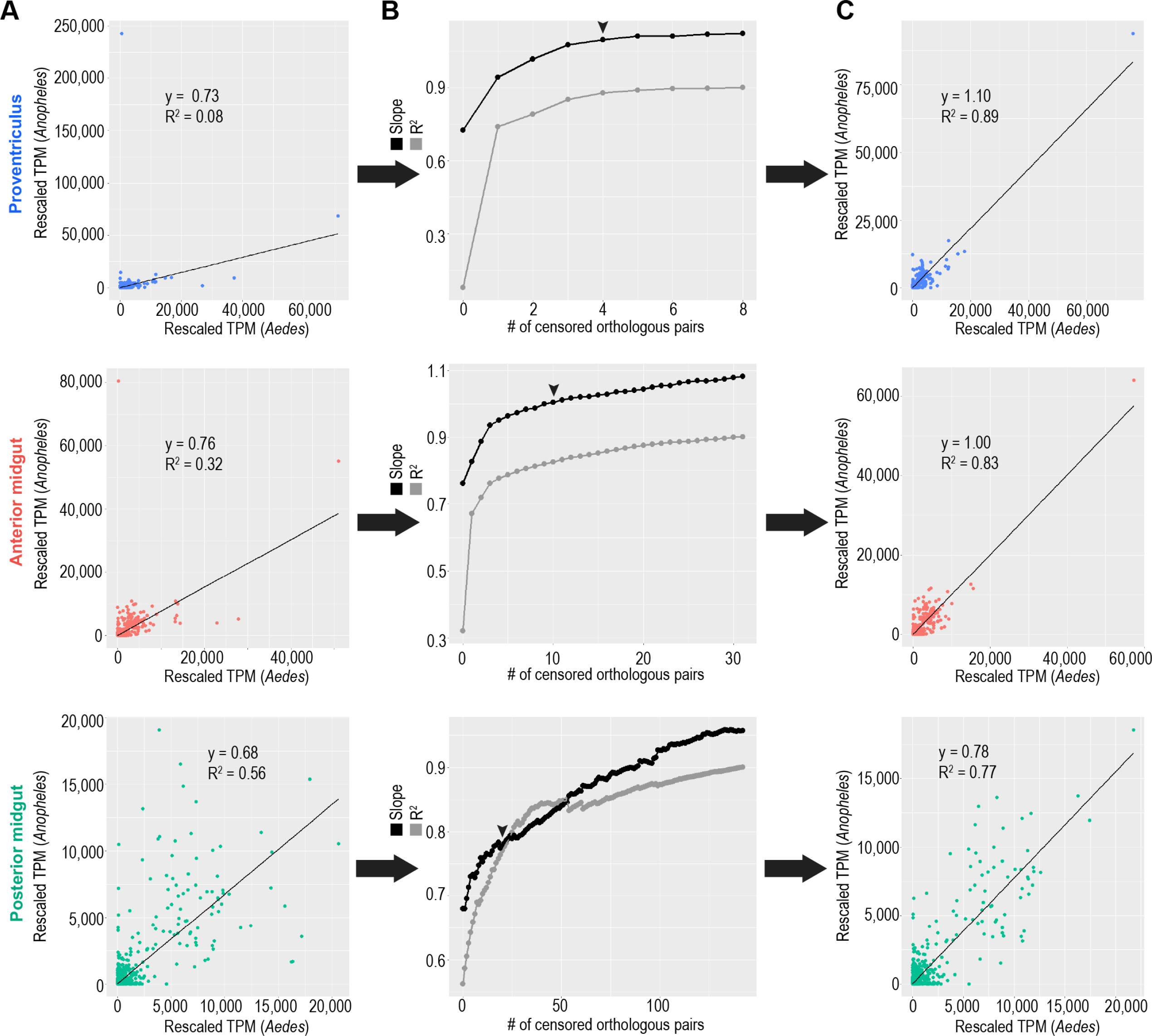
A small number of highly/disparately transcribed genes obscure the transcriptional correlation between one-to-one orthologous genes in *Aedes aegypti* and *Anopheles gambiae* midgut regions. (A) Correlation of expression of one-to-one orthologous genes in the *Aedes aegypti* and *Anopheles gambiae* midgut regions. (B) Plotted slopes and correlation coefficients for correlations of one-to-one orthologous genes during a process of sequential censoring of disparately expressed orthologs and re-scaling of TPMs; Black arrows indicate the number of genes censored in: (C) Adjusted correlation plots reflecting the relationships of one-to-one orthologs post-censorship/rescaling.

**S7.2 Fig.**
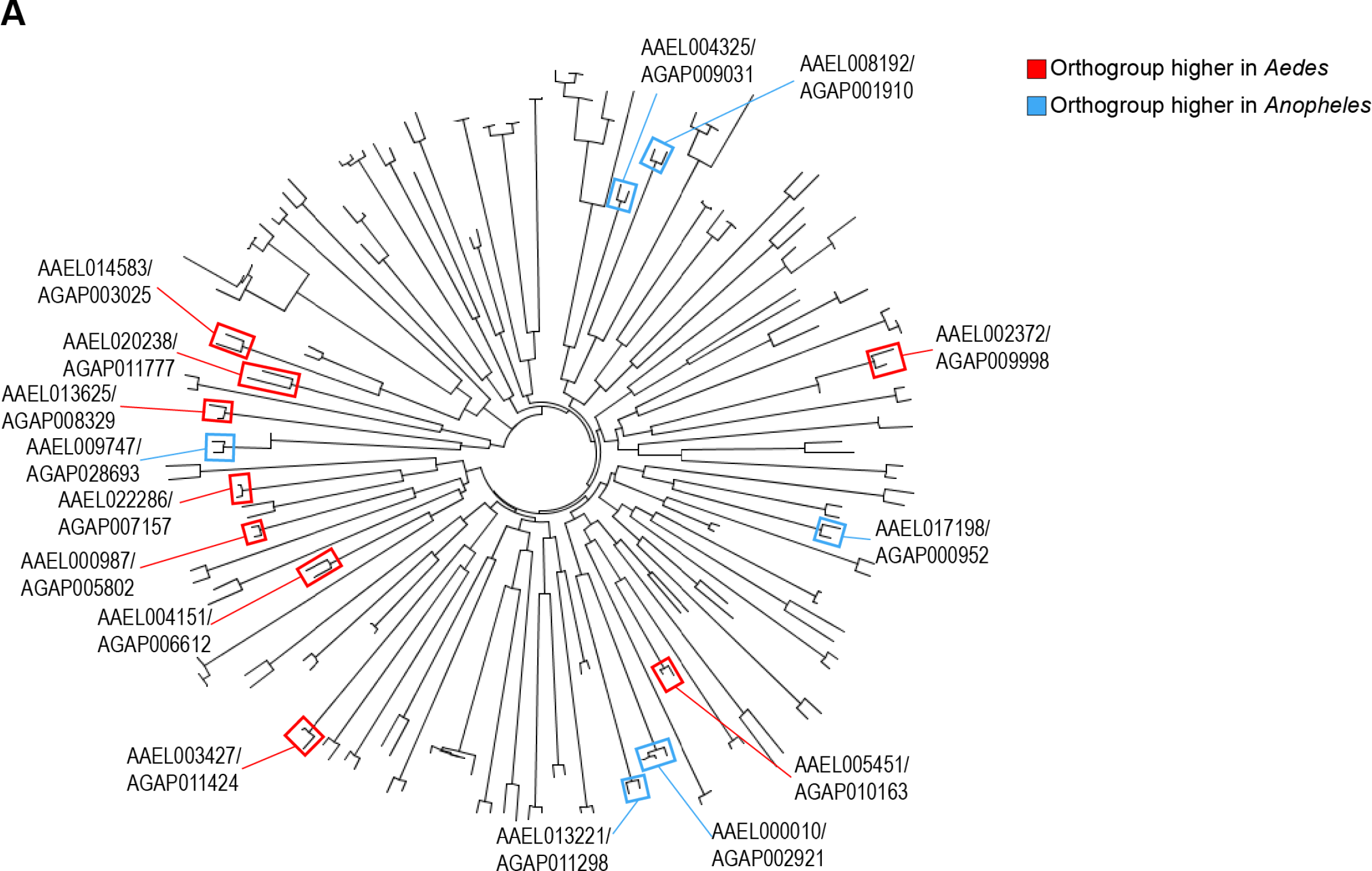
Phylogenetic analysis confirms disparately expressed one-to-one orthologous ribosomal proteins are sister sequences. (A) Geneious phylogenetic tree depicting the evolutionary relatedness of ribosomal proteins in *Aedes aegypti* and *Anopheles gambiae*. Pairs of censored one-to-one orthologs from the correlational analysis are shown in colored boxes. A red box indicates that the *Aedes* gene was higher expressed compared to the *Anopheles* gene. A blue box indicates that the *Anopheles* gene was higher expressed.

**S7.3 Fig.**
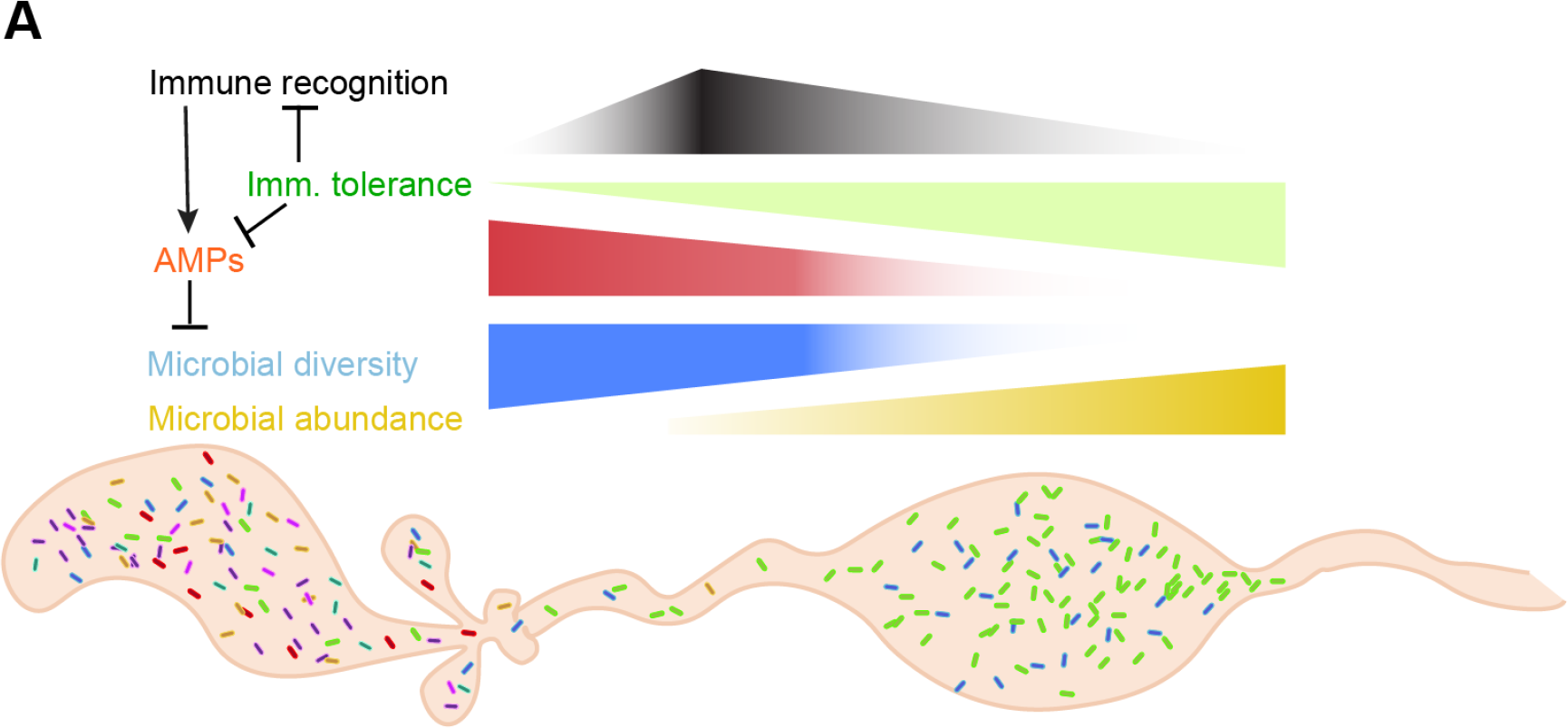
A model for regionalized microbe selection and immune tolerance in the mosquito gut. (A) A model of the proposed effects of regional specializations in immune recognition/AMP expression (proventriculus/anterior midgut) and tolerance (posterior midgut).

